# Peristromal niches protect lung cancers from targeted therapies through a combined effect of multiple molecular mediators

**DOI:** 10.1101/2024.04.24.590626

**Authors:** Bina Desai, Tatiana Miti, Sandhya Prabhakaran, Daria Miroshnychenko, Menkara Henry, Viktoriya Marusyk, Chandler Gatenbee, Marylin Bui, Jacob Scott, Philipp M. Altrock, Eric Haura, Alexander R.A. Anderson, David Basanta, Andriy Marusyk

**Affiliations:** Department of Metabolism and Physiology, H Lee Moffitt Cancer Centre and Research Institute, Tampa, FL, USA; Cancer Biology Ph.D. Program, University of South Florida, Tampa, FL; Department of Integrated Mathematical Oncology, H Lee Moffitt Cancer Center and Research Institute, Tampa, Florida; Department of Pathology, H. Lee Moffitt Cancer Center and Research Institute, Tampa, Florida; Department of Translational Hematology and Oncology Research, Cleveland Clinic, Cleveland, OH, USA; Department of Theoretical Biology, Max Planck Institute for Evolutionary Biology, Ploen, Germany; Department of Thoracic Oncology, H. Lee Moffitt Cancer Center and Research Institute, Tampa, FL, USA; Department of Molecular Medicine, University of South Florida, Tampa, Florida

## Abstract

Targeted therapies directed against oncogenic signaling addictions, such as inhibitors of ALK in ALK+ NSCLC often induce strong and durable clinical responses. However, they are not curative in metastatic cancers, as some tumor cells persist through therapy, eventually developing resistance. Therapy sensitivity can reflect not only cell-intrinsic mechanisms but also inputs from stromal microenvironment. Yet, the contribution of tumor stroma to therapeutic responses *in vivo* remains poorly defined. To address this gap of knowledge, we assessed the contribution of stroma-mediated resistance to therapeutic responses to the frontline ALK inhibitor alectinib in xenograft models of ALK+ NSCLC. We found that stroma-proximal tumor cells are partially protected against cytostatic effects of alectinib. This effect is observed not only in remission, but also during relapse, indicating the strong contribution of stroma-mediated resistance to both persistence and resistance. This therapy-protective effect of the stromal niche reflects a combined action of multiple mechanisms, including growth factors and extracellular matrix components. Consequently, despite improving alectinib responses, suppression of any individual resistance mechanism was insufficient to fully overcome the protective effect of stroma. Focusing on shared collateral sensitivity of persisters offered a superior therapeutic benefit, especially when using an antibody-drug conjugate with bystander effect to limit therapeutic escape. These findings indicate that stroma-mediated resistance might be the major contributor to both residual and progressing disease and highlight the limitation of focusing on suppressing a single resistance mechanism at a time.

## INTRODUCTION

In many cancers targeted therapies directed against oncogenic signaling addictions induce strong and durable responses that can keep tumors in remission for years, with minor side effects. Response of non-small cell lung cancers (NSCLC) with oncogenic EML4-ALK fusions to ALK inhibitors (ALKi) can be viewed as a “poster child” of this efficiency. Unfortunately, targeted therapies are not curative in metastatic cancers. A subset of tumor cells avoids elimination (persists) within residual tumors, and these residual populations eventually acquire net positive growth (*bona fide* resistance) responsible for tumor relapse^1^. Historically, both persistence and resistance have been attributed to cell-intrinsic mechanisms that define therapy sensitivity. However, growing evidence indicates that interactions with tumor stroma can strongly reduce therapeutic sensitivity^2–4^. This effect, which can be termed “environment-mediated drug resistance” (EMDR),^2^ has been described across a wide range of therapeutic contexts and appears to be particularly relevant to targeted therapy responses. Cancer-associated fibroblasts (CAFs), the main cellular component of tumor stroma, have emerged as a central mediator of EMDR^2, 4, 5^. Co-cultures of tumor cells with CAFs or CAF-conditioned media (CM) can significantly reduce their sensitivity to targeted therapies, including reduced sensitivity of ALK+ NSCLC cells to ALKi^6–8^. This effect can be mediated by CAF-produced cytokines, extracellular matrix (ECM) components, metabolites, and exosomal microRNAs^4, 9, 10^.

In contrast to the advanced understanding of the mechanistic underpinning of EMDR, its contribution to therapeutic responses *in vivo* remains poorly understood. Across a wide range of *in vitro* experimental models, tumor cells demonstrate cell intrinsic persistence^11^. Therefore, maintenance of residual disease can be expected to reflect a combination of EMDR and cell-intrinsic persistence. The potential contribution of EMDR to the *bona fide* relapse-driving resistance is less intuitively clear, as EMDR is mediated by paracrine-acting mechanisms. Therefore, EMDR-driven relapse would require a coordinated expansion of tumor cells and their protective niches,

To address the relevance of CAF-mediated EMDR to therapeutic responses *in vivo*, we decided to use a common model of ALK+ NSCLC, the H3122 cell line. Under standard 2D *in vitro* culture conditions devoid of stroma, subsets of H3122 cells persists the exposure to clinically relevant ALKi concentrations, gradually acquiring *bona fide* resistance^12^. At the same time, the *in vitro* sensitivity of H3122 cells to ALKi can be strongly reduced in the presence of CAFs. Therefore, *in vivo* responses of H3122 tumors to ALKi can be expected to include both cell-intrinsic and EMDR components. Given the multifactorial nature of adaptation of H3122 cells to ALKi ^12^, it is challenging to switch off cell-intrinsic persistence/resistance experimentally. Therefore, we attempted to dissect a relative contribution of EMDR to *in vivo* ALKi responses by therapeutic manipulation of a mechanism(s) that mediate the protective effect of CAFs.

Consistent with prior reports^7, 13^, we found that the ability of multiple primary CAF isolates to protect H3122 cells from ALKi *in vitro* is largely reducible to the activation of cMET by CAF-secreted hepatocyte growth factor (HGF). Surprisingly, modulation of the HGF-cMET axis in mouse xenograft models had only a modest impact on ALKi sensitivity of H3122 tumors. At the same time, our spatial analyses indicated a strong contribution of EMDR to both persistence and resistance in experimental mouse tumors. We found that, like the multifactorial nature of cell-intrinsic resistance^12, 14^, EMDR cannot be reduced to a single mechanism. Instead, it combines the effects of multiple growth factors and ECM components. Consequently, despite enhancing tumor regression and delaying relapse, pharmacological suppression of individual mechanisms could not fully overcome EMDR effects. To bypass the challenge posed by the multifactorial nature of therapy persistence and resistance, we explored the utility of targeting a strong collateral sensitivity to EGFR/HER2 inhibition shared between stroma-mediated and cell-intrinsic persistence. Whereas a dual EGFR/HER2 inhibitor lapatinib strongly enhanced alectinib sensitivity and extended remission, residual tumors eventually developed resistance to the combined therapy and relapsed. To further extend therapeutic control, we explored the utility of combining alectinib with T-DXd, an antibody-drug conjugate that combines targeting HER2+ expressing cells with suppression of HER2-bystanders. Alectinib-T-DXd combination induced the strongest regression, eradicating some of the residual tumors. Our results indicate that spatially limited EMDR might be a major mediator of residual disease and an enhancer of tumor relapse. Further, our findings highlight the limitations of focusing on individual resistance mechanisms. A more promising approach is to focus on shared collateral sensitivities and broadening therapeutic responses by eliciting bystander effects.

## RESULTS

### Stromal fibroblasts reduce the in vitro sensitivity of NSCLC cells to targeted therapies via HGF-cMET axis

As the first step to deciphering the impact of EMDR on *in vivo* responses of the H3122 model of ALK+ NSCLC to ALKi, we sought to define a mechanism (s), responsible for the ability of CAFs to reduce the sensitivity of H3122 cells to ALKi^8, 15–17^. Consistent with our prior report^15^, co-culture with a primary lung cancer CAF isolate (subsequently referred to as CAF1) substantially reduced the sensitivity of H3122 to the frontline ALKi alectinib (**Fig. 1A**). CAF1 CM fully phenocopied the effect of a physical CAF1 co-culture, indicating that the protective effect is mediated by a secreted factor(s) (**Fig. 1A**). CAF1 CM conferred similarly strong protection to clinically approved ALKi, lorlatinib, ceritinib and brigatinib, but not to the dual ALK and cMET inhibitor crizotinib (**Fig. 1B**). A similar effect was observed in the ALK+ NSCLC STE-1 cell line (**Fig. S1A**). Moreover, consistent with prior reports, CAF1 CM conferred a strong protection against EGFR inhibitors gefitinib and erlotinib in the PC9 cell line model of EGFR mutant NSCLC (**Fig. S1B**) as well as against KRAS^G12C^ inhibitors sotorasib and ARS-1620 in the H358 model of KRAS^G12C^ NSCLC (**Fig. S1C)**. Next, given the known phenotypic diversity of fibroblasts^8, 18^, we examined the effect CM from a panel of primary human NSCLC CAF isolates as well as fibroblasts isolated from normal lung tissue (NF) (**Fig. S1D**). Despite a variability in the magnitude of the effect, CM from most fibroblast isolates protected H3122 cells against alectinib, lorlatinib, ceritinib, and brigatinib, while having little to no protection against crizotinib (**Fig. 1C**).

**Fig. 1.**
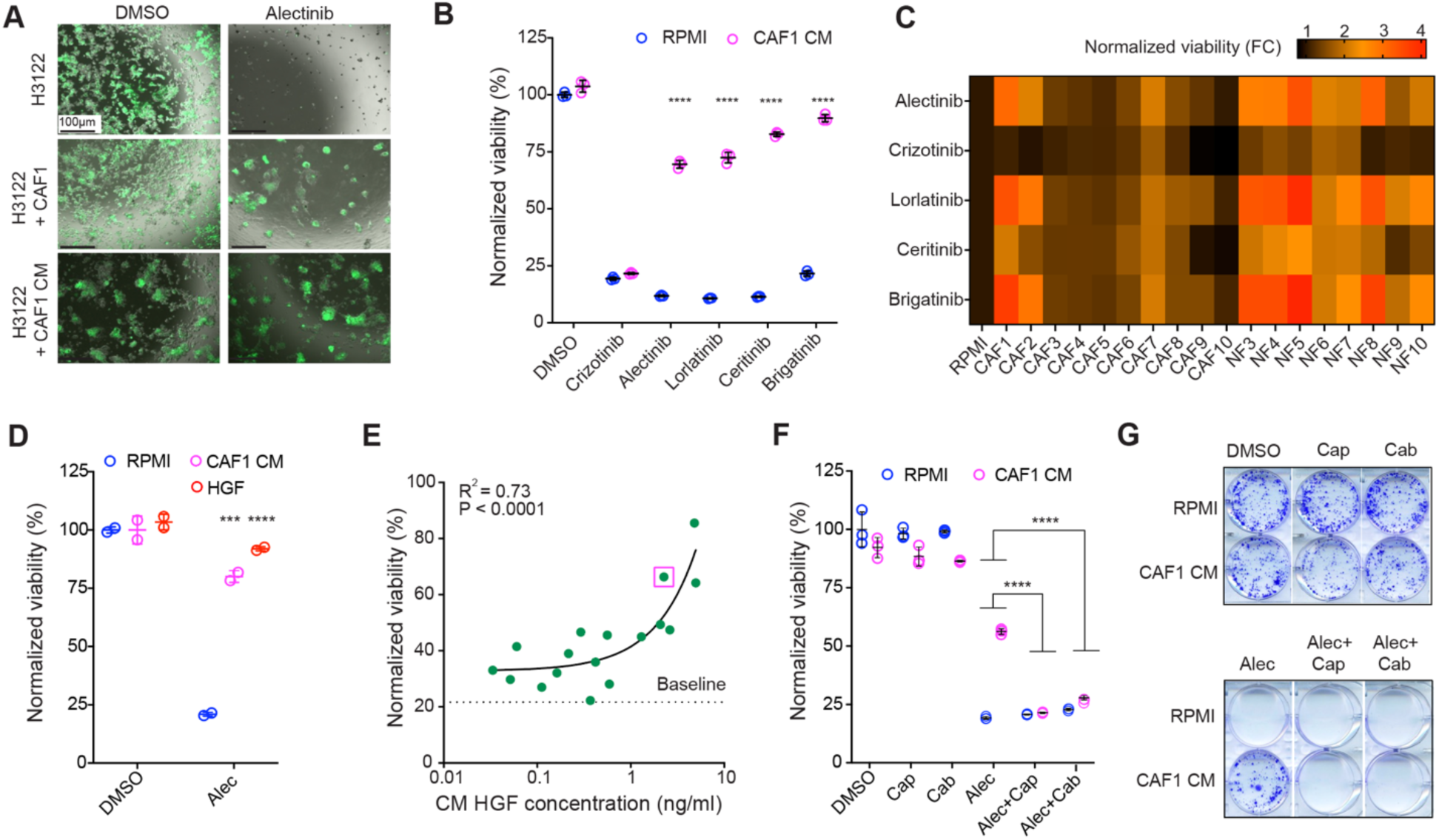
Fibroblasts protect ALK+ NSCLC from ALKi through the HGF-cMET axis. **A**. Live fluorescent microscopy images of GFP labeled H3122 cells co-cultured with unlabeled CAFs or CAF1 CM in the presence of alectinib (0.5μM) or DMSO vehicle control. **B**. Cell Titer-Glo viability assay of H3122 cells cultured in RPMI or CAF CM with DMSO control, crizotinib (0.5 µM), alectinib (0.25 µM), lorlatinib (0.5 µM), ceritinib (0.1 µM) and brigatinib (0.25 µM). Results are normalized to the DMSO control signal in RPMI. **C**. Heatmap summary of the impact of fibroblast CM on sensitivity to H3122 cells to the indicated ALKi. The sensitivity is presented as the fold change in cell viability as compared to the RMPI media control**. D**. Exogenous HGF at the concentration found in CAF1 CM (2.25 ng/ml) phenocopies alectinib-protecting effect of CAF1 CM. **E**. Pearson correlation analysis of the correlation between HGF levels detected in the CM of individual fibroblast isolates and the impact of CM on alectinib (0.1 µM) sensitivity. Purple square highlights CAF1 CM. **F, G**. cMET inhibitors capmatinib (0.2 µM) and cabozantinib (0.2 µM) overcome protective effect of CAF1 CM to alectinib (0.25 µM) in short-term (4d) viability assay (F) and longer-term clonogenic survival assay (**10d**), (**G**). *** and **** indicate p<0.001 and p<0.0001, respectively, of the interaction term of two-way ANOVA assay, comparing the impact of CM on viability of H3122 cells under treatment with ALKi versus vehicle control (DMSO) (**B, D**), or on viability of H3122 cells under alectinib in the absence or presence of cMET inhibitors (**F**).

The lack of protection against crizotinib suggested mediation by the well-established bypass signaling, where stromal HGF activates its unique receptor cMET expressed by tumor cells ^7, 17, 19^. Indeed, exogenous HGF conferred strong dose-dependent protection against alectinib in H3122 and STE-1 models (**Fig. S2A, B**). Moreover, exogenous HGF, added at the concentrations detected in CAF1 CM (2 ng/ml) fully phenocopied the protective effect of CAF1 CM against alectinib and lorlatinib (**Fig. 1D, S2C**), while providing no detectable protection against crizotinib (**Fig. S2C**). Importantly, we observed a strong correlation (R^2^=0.73) between HGF concentration in the CM of different fibroblast isolates and the magnitude of their protection of H3122 against alectinib (**Fig 1E**), indicating a generalizability of the CAF1 CM inferences. Consistent with the central role of HGF-cMET axis in mediating the effects of CAFs, the addition of HGF-neutralizing antibody strongly suppressed the protective effects of CAF CM in H3122 and STE-1 models (**Fig. S2D, E**). Similarly, cMET inhibitors cabozantinib and capmatinib completely suppressed CAF-mediated protection in our experimental models of ALK+, EGFR mutant and KRAS^G12C^ NSCLC (**Fig. 1F, S2F-I**). Importantly, protection observed within short-term growth assay translated into longer-term protection observed in clonogenic assays (**Fig. 1G, Fig. S2J**). In summary, the substantial reduction of ALKi sensitivity of H3122 cells in the presence of fibroblast CM reflects a general phenomenon of CAF-mediated targeted therapy resistance. At least *in vitro*, the protective effect of CAFs in the H3122 model can be largely reduced to the well-characterized HGF-cMET bypass signaling mechanism.

### The HGF-cMET axis moderately impacts alectinib responses in vivo

Assuming the conservation of the mechanisms mediating the effect of CAFs between *in vitro* and *in vivo* contexts, the contribution of EMDR to alectinib responses of H3122 tumors *in vivo* could be assessed by experimental manipulation of the HGF-cMET axis in xenograft models. However, murine HGF does not activate human cMET^20^ (**Fig. S2A**), complicating the assessment as EMDR might be irrelevant in conventional xenograft models.

To overcome this limitation, we first evaluated the ability of secreted HGF to mitigate alectinib sensitivity *in vivo*. To this end, we overexpressed human HGF in H3122 cells using a lentiviral vector and compared alectinib responses between NSG xenograft tumors formed by subcutaneous grafts of H3122 parental cells (which lack detectable HGF expression) and their HGF overexpressing counterparts. Whereas HGF expression had no discernable impact on the baseline tumor growth, it enabled tumors to grow under alectinib treatment that induced strong regression of the parental H3122 tumors (**Fig. 2A, S3B**). However, the ability of HGF to provide a *bona fide* alectinib resistance in the H3122 model *in vivo* does not fully address the impact of stromal HGF, as the effects of paracrine-acting HGF can be spatially restricted to stroma proximal niches. To evaluate the impact of the HGF-mediated EMDR more directly, we leveraged the compatibility between human HGF and murine cMET, as a replacement of both murine HGF alleles in the NSG mice with their human counterparts produces no overt phenotype, apart from modulating responses of HGF-responsive human xenografts^21^. Thus, analyses of therapy responsiveness of parental H3122 cells grafted in the NSG strain versus its humanized derivate (NSG^hHGF^) ensure for a “clean” evaluation of the impact of stromal HGF on therapeutic responses (**Fig. 2B**).

**Fig. 2.**
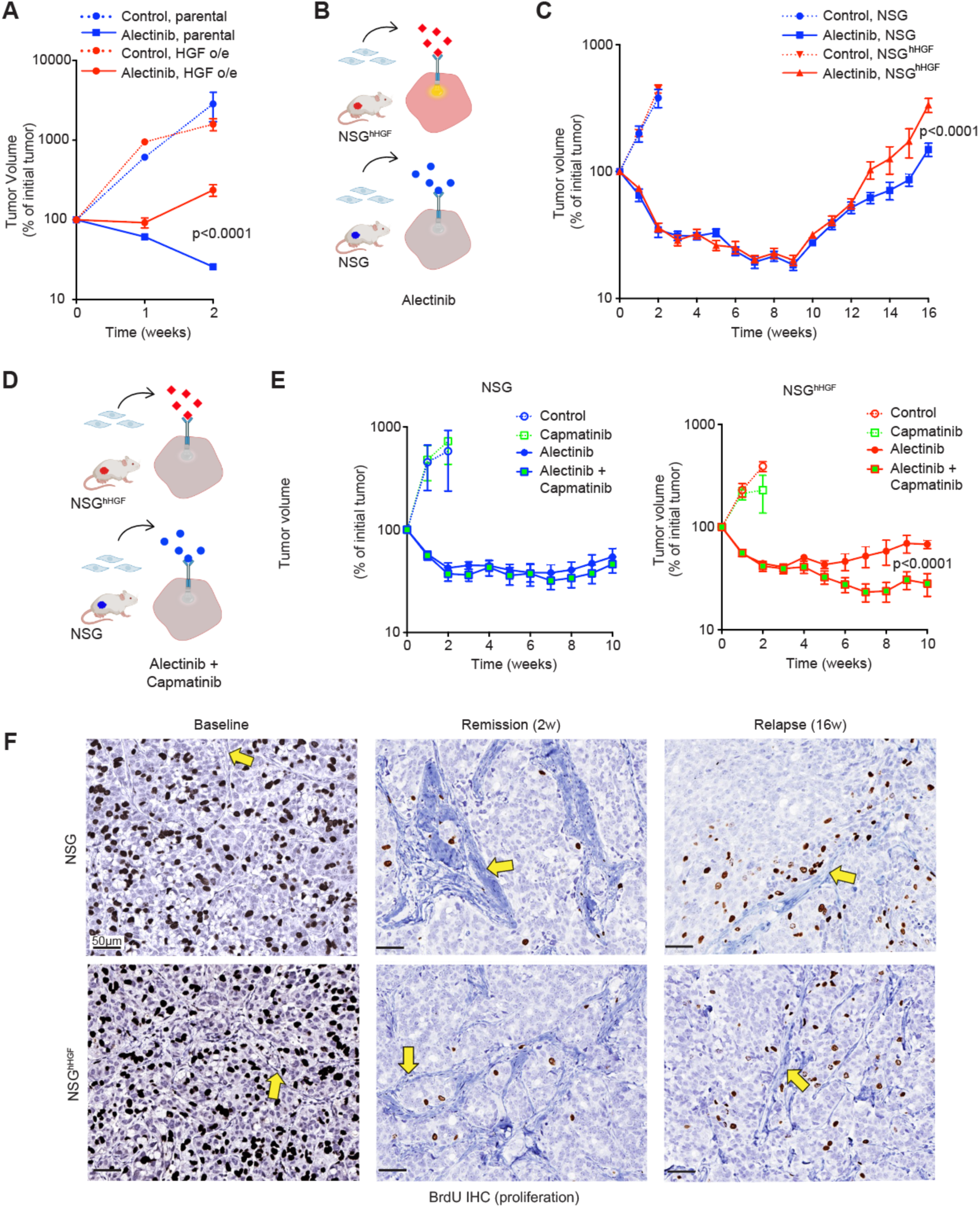
Xenograft studies indicate HGF-independent protection of H3122 cells from alectinib in peristromal niches. **A.** Exogenous expression of HGF confers resistance to alectinib (25 mg/kg) in NSG xenografts. N=4 tumors for all groups except for HGF o/e control (N=2). Error bars represent SEM. **B**. Diagram of the experimental design to assess the impact of stromal activation of tumor cMET -mediated bypass signaling *in vivo*. **C**. Volumetric response dynamics of H3122 xenograft tumors in NSG and NSG^hHGF^ hosts, continuously treated with 25 mg/kg alectinib or vehicle control. Error bars represent SEM; N= 4 & 18 for the control and alectinib-treated groups, respectively. **D**. Diagram of the experimental idea: cMET inhibition should suppress the enhanced relapse in NSG^hHGF^ xenografts. **E**. Tumor response dynamics for vehicle control (N=4), 40 mg/kg capmatinib (N=4), 20 mg/kg alectinib (N=10) and alectinib/capmatinib (N=10) combination treated groups respectively. **F**. Representative images of anti-BrdU IHC from xenograft tumor tissues at the indicated time points. Arrows point to the examples of stromal regions defined by counterstain. P values in tumor growth analyses represents the significance of the interaction term of the repeated measurement 2-way ANOVA analyses between the indicated groups.

Consistent with the lack of impact of HGF on the baseline proliferation of H3122 cells both *in vitro* (**Fig. 1D**) and *in vivo* (**Fig. 2A**), we observed no discernable differences in tumor implantation and growth before the initiation of the treatment (**Fig. S3C**) or under vehicle control (**Fig. 2C, S3D**). Surprisingly, stromal expression of human HGF did not impact the initial tumor responses, and the reduced alectinib sensitivity of NSG^hHGF^ grafts was only detectable at relapse (**Fig. 2C, S3D, E**). Given the relatively modest effect of humanized stromal HGF, we further validated the impact of the HGF-cMET axis using a highly specific cMET inhibitor, capmatinib (**Fig. 2D**). To enhance the ability to resolve modest differences in alectinib sensitivity, we used a slightly lower alectinib dosing (20 mg/kg instead of 25 mg/kg). Consistent with the inability of murine HGF to activate human cMET, capmatinib had no discernable effect on alectinib response of H3122 tumors grafted into NSG mice. In contrast, in NSG^hHGF^ H3122 xenografts, capmatinib substantially enhanced the magnitude of tumor regression and delayed the relapse (**Fig. 2E, S3F, G**), supporting the relevance of the HGF-cMET axis for stroma-mediated resistance *in vivo*.

### Proximity to stroma reduces alectinib sensitivity in vivo through both-dependent and HGF-independent mechanisms

Given the strong effect of ectopic HGF expression *in vivo* (**Fig. 2A**) and the ability of fibroblast-produced HGF to provide ALKi protection *in vitro* (**Fig. 1**), we reasoned that the modest effect of stromal HGF in xenograft models could reflect the spatial limitation of cMET activation to tumor cells in the immediate proximity to HGF-producing stroma. To address this possibility, we examined the spatial distribution of tumor cell proliferation in the two xenograft models, using IHC staining for a BrdU which marks cells undergoing DNA replication. Without treatment, the distribution of BrdU+ cells appeared to be random in both the NSG and NSG^hHGF^ xenograft tumors (**Fig. 2F**, baseline). Consistent with therapy-protecting effects of CAFs *in vitro*, alectinib treatment potently suppressed cell proliferation while shifting the distribution of BrdU+ cells to peristromal niches (**Fig. 2F**, remission). Unexpectedly, this treatment-induced bias was observed in both xenograft tumor models, suggesting the existence of a strong HGF-independent component of EMDR (**Fig. 2F**). Intriguingly, the bias in BrdU+ staining towards stroma-proximal spatial niches was not limited to remission. As expected, tumor growth relapse was associated with a marked increase in tumor cell proliferation and the expansion of tumor parenchyma, indicating an increase in cell-intrinsic resistance. At the same time, the bias in spatial distribution of BrdU+ cells toward stroma-proximal niches was partially retained in relapsing tumors in both xenograft models (**Fig. 2F**, relapse). This retention of the spatial bias indicated that the stroma-mediated resistance contributes to the net positive tumor cell growth that drives the relapse.

To quantitatively assess the impact of stroma proximity on tumor cell proliferation and to evaluate the contribution of the HGF-cMET axis, we took advantage of our recently published digital pathology/spatial analysis pipeline^22^. In this approach, scanned microscopy images of the anti-BrdU IHC staining of histological tumor cross-sections are segmented into necrotic areas (excluded from the analyses), stromal regions, BrdU+ and BrdU-tumor cells. Continuous stromal regions, defined by clearly distinguishable ECM deposition, are further rasterized to facilitate quantitation.

Our ability to draw accurate inferences from spatial distribution requires adequate controls. Since therapy reduces the tumor/stroma ratio and alters the relative distribution of tumor parenchyma and stroma, direct comparisons between the control and alectinib-treated tumors are not easily interpretable. To overcome this limitation, the spatial distributions of proliferating BrdU+ tumor cells are compared to their digital sample-specific digital “avatars”. These digital “avatars” retain specific spatial locations of each tumor cell and stromal regions, but the BrdU positivity status is randomly redistributed among the tumor cells while preserving the BrdU+ fraction (**Fig. 3A**). Comparison between the observed and predicted random distributions enable accurate assessment of the spatial proliferation bias in each sample. Comparisons of pairwise (observed versus predicted) differences between vehicle control and alectinib-treated groups enable the assessment of the impact of therapy. Finally, comparisons of the predicted/observed differences between the NSG and NSG^hHGF^ xenograft tumors enable the assessment of the impact of stromal HGF. Importantly, our pipeline analyses all tumor cells within the histological cross-sections (80K-400K per tumor), with multiple independent tumors analyzed per time point, thus minimizing the impacts of spatial heterogeneity and sampling biases.

**Fig. 3.**
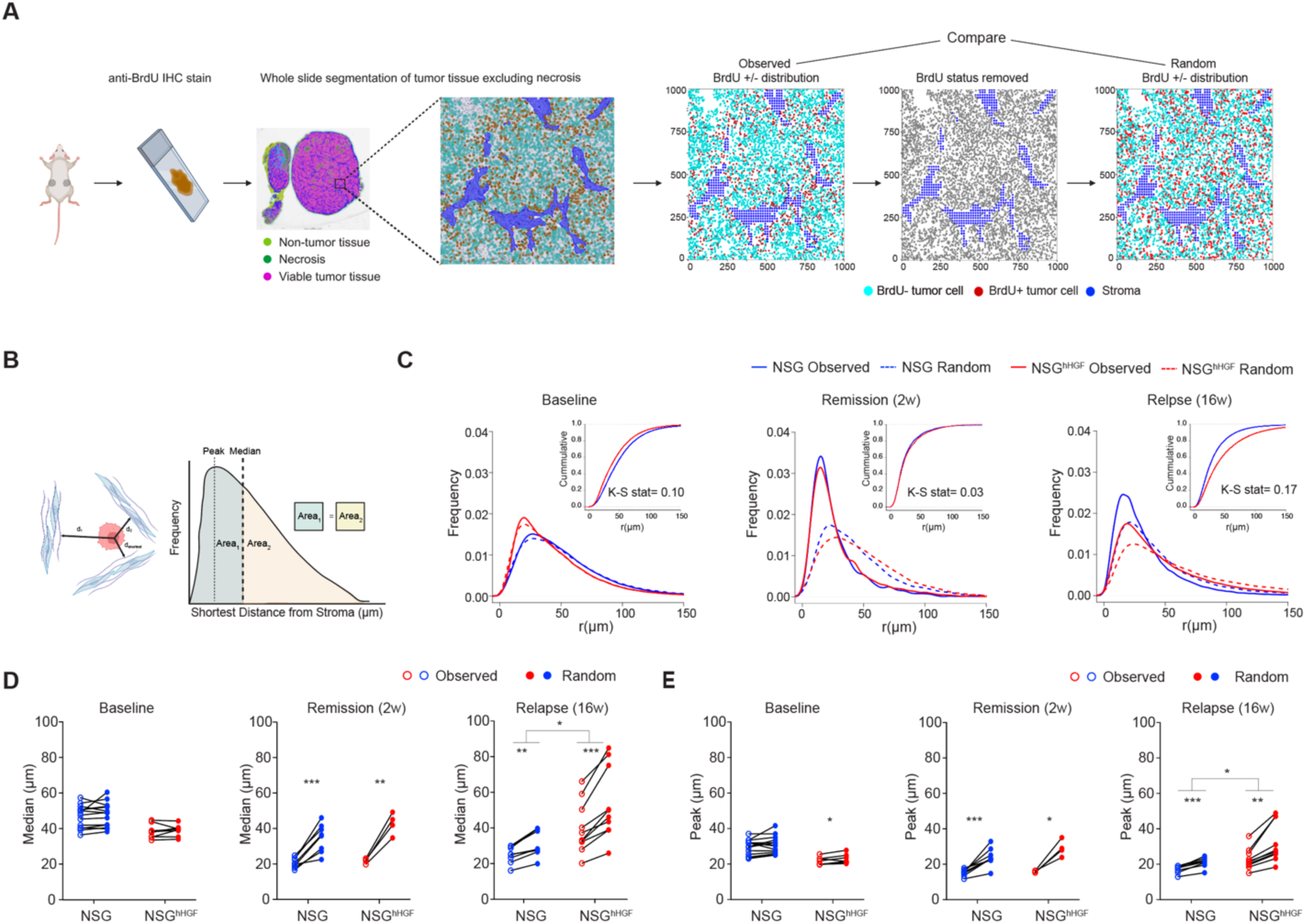
Spatial analyses of tumor tissues indicate HGF-independent sheltering from cytostatic effects of alectinib in peristromal niches. **A.** Diagram of the spatial analysis workflow. **B.** Diagram depicting analyses of distances to the nearest stroma. **C.** Observed and random distributions of the distances of BrdU+ cells to the nearest stroma; cumulative distribution and KS statistics values are shown as insets. **D, E.** Ladder plots comparing the medians **(D)** and means **(E)** of predicted and observed distributions of the distances of BrdU+ cells to the nearest stroma. Each point reflects an analysis of a whole cross-section of an individual tumor. *, **, *** indicate p values of less than 0.05, 0.01, and 0.001, respectively. Comparisons between the observed and predicted values were performed using a 2-tailed t-test; comparisons between NSG and NSG^hHGF^ models were performed with the analyses of the interaction term of 2-way ANOVA test. KS denotes Kolmogorov-Smirnov test.

To assess the impact of stroma proximity on cell proliferation, we assessed the distributions of distances of BrdU+ tumor cells to the nearest stroma^23, 24^ (**Fig. 3B**), with the expected random distributions serving as sample-specific controls. Our analyses confirmed the visual inferences of the lack of the spatial bias in NSG and NSG^hHGF^ xenograft tumors in the absence of therapy (**Fig. 3C-E**). In contrast, we detected a strong spatial bias in BrdU+ tumor cells towards peri-stromal regions in alectinib treated tumors. While this bias was most pronounced in regressing tumors, it was also present in relapsing tumors (**Fig. 3C-E**). Consistent with the lack of detectable differences in the tumor regression (**Fig. 2D**), we did not detect differences in the stroma proximity bias between the NSG and NSG^hHGF^ tumors during remission. However, at relapse, the bias was significantly larger in the NSG^hHGF^ xenografts (**Fig. 3C-E**), consistent with our observation from Fig.2C. Taken together, our spatial analyses support the notion that stroma-mediated resistance contributes to both incomplete tumor responses and tumor growth relapse, and indicate that, *in vivo*, stroma-mediated resistance cannot be reduced to the effects of the HGF-cMET axis.

To uncover the potential source(s) of the HGF-independent EMDR component, we performed transcriptomic analyses of treatment-naïve and alectinib-treated (2 and 6 weeks) tumors from the H3122 NSG and H3122 NSG^hHGF^ xenograft models using bulk Illumina RNA sequencing. Reads were mapped to both human and murine genomes, enabling simultaneous assessment of neoplastic and stromal compartments (**Fig. 4A**). Principle component analysis (PCA) and non-supervised hierarchical clustering of the neoplastic transcriptomes clearly distinguished samples from treatment naïve and alectinib-treated tumors at different treatment timepoints (**Fig. 4B, Fig S4A**). Consistent with the lack of the baseline and early response differences between the NSG^hHGF^ and NSG xenografts in tumor growth and spatial distributions of cell proliferation, transcriptional analyses revealed the emergence of the HGF-dependent differences only after six weeks of treatment (**Fig. 4B, Fig. S4A**). Pathway enrichment analyses indicated therapy-induced, HGF-independent suppression of multiple pathways related to cell cycle, metabolism, and signaling, as well as therapy-induced activation of PI3-AKT signaling, cellular senescence, and ECM-receptor interactions (**Fig 4C**). To assess the utility of transcriptional inferences in identifying potential therapeutic vulnerabilities, we assessed the impact of suppressing PI3K signaling, one of the pathways differentially expressed between therapy-naïve and alectinib treated tumors with pharmacological inhibitor alpelisib. Consistent with a recent report^25^, alpelisib treatment significantly enhanced tumor responses of xenograft tumors to alectinib, supporting the utility of the transcriptomics inferences in identifying pathways mediating alectinib resistance (**Fig. S4B**).

**Figure 4.**
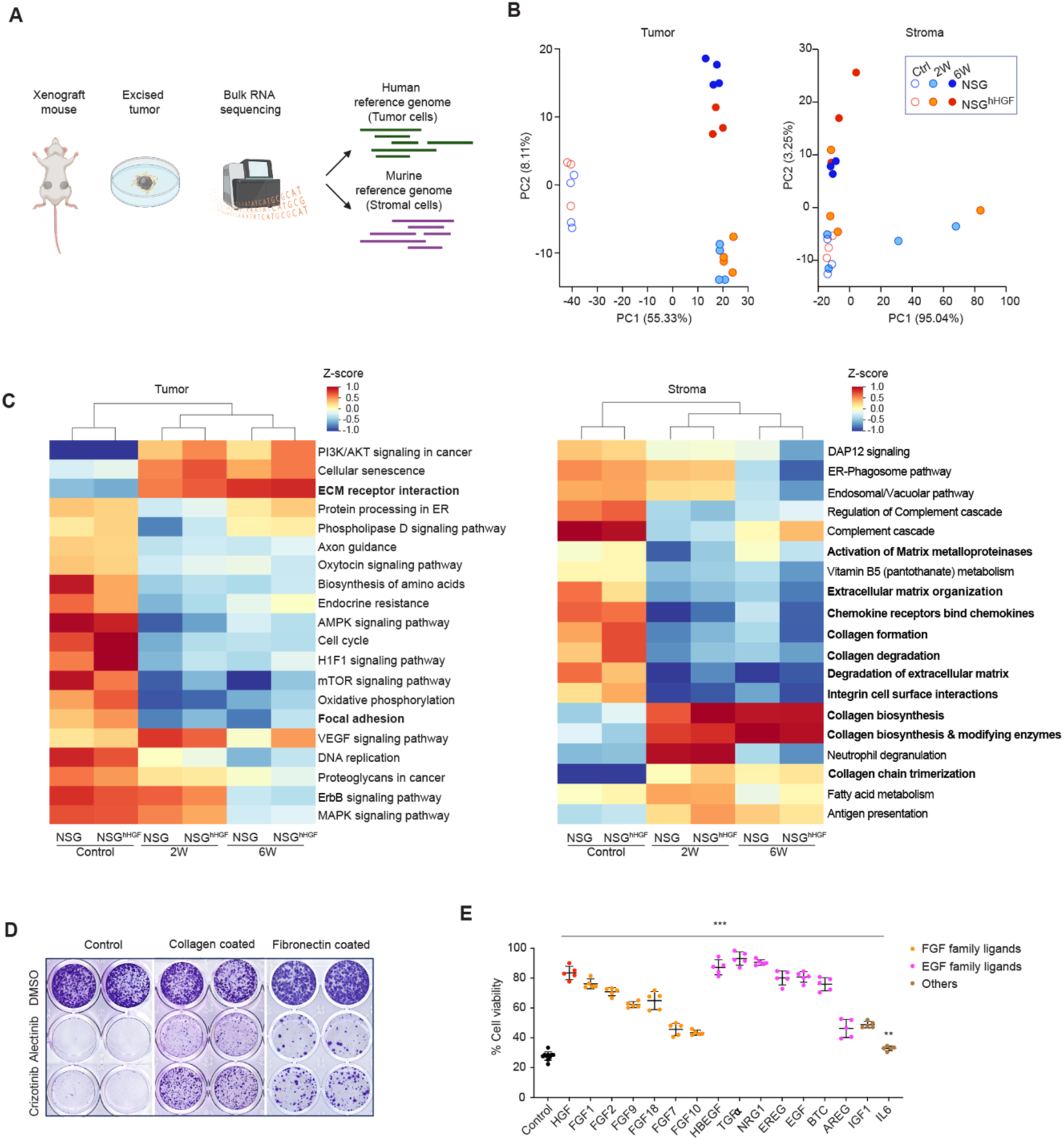
Transcriptomic analyses of tumor and stromal components identify the potential HGF-independent mechanisms of stroma-mediated therapy resistance. **A**. Diagram for transcriptional analyses approach. **B**. PCA analyses of neoplastic and stromal transcriptomes. **C**. Non-supervised hierarchical clustering of pathway analyses of differentially expressed genes. Bolded font indicates pathways related to ECM production and remodeling. **D**. Crystal violet stain of clonogenic assay of H3122 cells seeded into control, collagen or fibronectin coated plates cultured for 20 days in 0.1 µM alectinib, 0.5 µM crizotinib or DMSO control. **E**. Cell Titer Glo viability assay for H3122 cells cultured in 0.1 µM alectinib in the presence of indicated growth factors (50ng/ml). ** and *** indicate p<0.01 and p<0.001 respectively for the 2-tailed Mann Whitney test comparing alectinib sensitivity between the control samples and samples treated with the indicated growth factors

PCA and hierarchical clustering analyses of transcriptional reads mapped to mouse genome revealed a clear impact of alectinib treatment on global transcriptomes. However, the effects of the treatment duration and stromal expression of human HGF were less pronounced (**Fig. 4 B, Fig. S4A**). Pathway enrichment analyses revealed marked changes in multiple pathways involved in ECM synthesis, degradation, and organization in alectinib-treated tumors (**Fig. 4C**). Consistent with these changes, Mason Trichrome staining of histological samples revealed a therapy-induced enhancement of collagen deposition (**Fig. S4C**). Additionally, IHC staining with hyaluronan-binding protein revealed a substantial increase in the deposition of hyaluronan, which was previously implied in microenvironmental therapy resistance^24, 26^ (**Fig. S4D**).

Even though ECM-tumor cell interaction was dispensable for the observed *in vitro* effect of CAFs (**Fig. 1A**), integrin-mediated cell adhesion to collagens and fibronectin is a well-established microenvironmental resistance mechanism documented in multiple contexts of targeted therapies^27, 28^. Murine stroma expressed multiple ECM components implied in microenvironmental resistance, while their expression was undetectable in human neoplastic cells (**Fig. S4E**). To assess the potential contribution of collagens and fibronectin on the therapeutic sensitivity of ALK+ NSCLC to ALKi, we contrasted the responses of H3122 and STE-1 cells grown on regular tissue culture plastic with those grown on collagen-coated or fibronectin-coated plates. We found that cells grown on the collagen and fibronectin matrix were substantially less sensitive to both alectinib and crizotinib (**Fig. 4D, S5A**), supporting the potential involvement of cell-ECM interactions in HGF-independent mediation of EMDR to alectinib.

Next, we evaluated the potential involvement of cytokines and growth factors previously implied in microenvironment-mediated resistance to targeted therapies^4, 9, 10^. expression of these growth factor genes was primarily confined to the stromal compartment (**Fig. S4E**). Importantly, while not all these potential mediators displayed a detectable effect on alectinib sensitivity *in vitro*, multiple EGF and FGF family growth factors as well as IGF1 and IL6, substantially reduced the sensitivity of H3122 cells to alectinib in short-term growth assays and clonogenic studies (**Fig. 4E, Fig S5B, C**), supporting their potential role in mediating the HGF-independent effects of tumor stroma on alectinib sensitivity in vivo. In summary, our analyses suggest the multifactorial mechanism of EMDR, reflecting an integrated contribution of multiple interactions, including adhesion to ECM and the paracrine effect of multiple stroma-produced growth factors.

### Suppression of individual mechanisms of EMDR enhances in vivo responses but fails to fully abrogate the protective impact of stromal niches

To assess the contribution of HGF-independent mechanisms of EMDR to the alectinib sensitivity of tumors *in vivo*, we evaluated the effect of pharmacological suppression of the candidate mechanisms on the alectinib sensitivity of tumors implanted in NSG mice, lacking the ability to activate human cMET. To assess the impact of ECM-mediated adhesion, we used pirfenidone, a well-established anti-fibrotic and anti-inflammatory agent that suppresses TGF-β production^29, 30^. Importantly, TGF-β has no direct effect on reducing the sensitivity of H3122 cells to alectinib (**Fig. S5B-C**). Pirfenidone co-administration reduced the alectinib-induced enhancement in the deposition of collagens in H3122 xenografts (**Fig. 5A**) and substantially enhanced both the magnitude and the duration of remission (**Fig. 5B, S6A**). A similar enhancement of alectinib sensitivity was observed with a co-inhibition of focal adhesion kinase (FAK), which integrates signaling induced by integrin-mediated adhesion to collagens and fibronectins. FAK inhibitor defactinib enhanced responses to alectinib (**Fig. 5C, S6B**). To assess the contribution of hyaluronan, we evaluated the effect of the PEGylated recombinant human hyaluronidase (PEGPH20)^31^, which substantially reduced alectinib-induced hyaluronan deposition **5A**). (PEGPH20)^31^, which substantially reduced alectinib-induced hyaluronan deposition (**Fig. 5A**). Similar to the effect of suppressing collagen-integrin interaction, hyaluronidase co-treatment substantially enhanced alectinib sensitivity of H3122 tumors in NSG xenografts (**Fig. 5B, S6A**).

**Figure 5.**
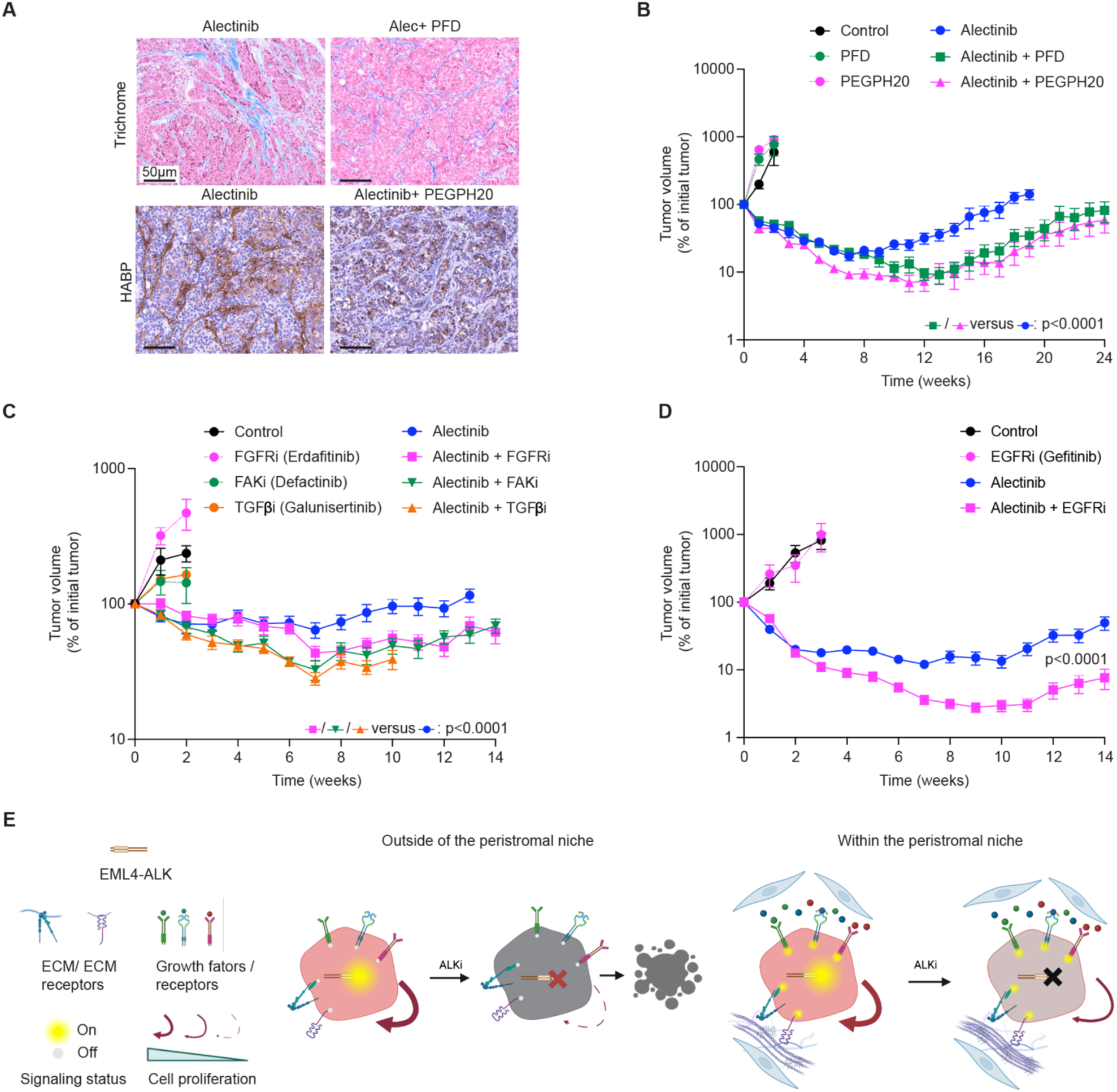
Suppression of multiple candidate mechanisms of stromal resistance improves responses to ALKi. **A**. Pirfenidone and PEGPH20 treatment suppresses alectinib-induced enhancement of collagen (Mason Trichrome staining, upper panel) and hyaluronan (lower panel) deposition, respectively. **B.** Volumetric responses of H3122 xenograft tumors to vehicle (N=4), 900 mg/kg pirfenidone (N=6), 0.1 mg/kg PEGPH20 (N=6), 25 mg/kg alectinib (N=10), alectinib/PEGPH20 and alectinib/pirfenidone combinations (N=10). **C**. Volumetric responses of H3122 xenograft tumors to vehicle, 20 mg/kg erdafitinib, 50 mg/kg defactinib, 75 mg/kg galunisertib, 20 mg/kg alectinib and the indicated combinations. N=6 for the control and individual co-inhibitors. N=10 for the alectinib and the combination treatment groups. **D**. Volumetric responses of H3122 xenograft tumors to vehicle, 20 mg/kg alectinib, 40 mg/kg gefitinib and alectinib/gefitinib combination. N=10 for the alectinib and combination treatment groups, N=6 for the control and gefitinib group. P values indicate the result of the interaction group of repeated measurement ANOVA comparing combination therapies to the alectinib monotherapy. **E**. Diagram of the conceptual model. Peristromal niche location enables tumor cells to partially maintain viability and proliferation upon ALK inhibition due to multiple ECM and growth factor signaling mechanisms. Outside of the peristromal niche, cells lack the compensating mechanisms, leading to a more complete proliferation arrest and apoptosis.

Next, we evaluated the relevance of stromal growth factors to the HGF-independent effects of stromal protection. To this end, we assessed the impact of FGFR and EGFR inhibition. Consistent with our *in vitro* data (**Fig. 4E, S5C**), FGFR inhibitors erdafitinib and infigratinib, and EGFR inhibitor gefitinib, enhanced alectinib sensitivity, delaying tumor growth remissions (**Fig. 5C, D, S6 B-D**). Consistent with incomplete effects of the tested therapy combinations, histological analyses of cell proliferation revealed that none of the inhibitors of individual EMDR mechanisms was able to completely overcome proliferation of tumor cells within peristromal niches (**Fig. S6E**). In summary, our results suggest that similar to the multifactorial nature of cell-intrinsic resistance mechanisms^14, 15^, EMDR cannot be reduced to a single molecular pathway. Instead, it integrates the functional impact of multiple mechanisms enabling tumor cells to escape therapeutic elimination within peristromal niches and contributing to tumor growth relapse (**Fig. 5E).**

### Targeting collateral sensitivities enables to bypass EMDR

Multifactorial nature of EMDR poses an obvious limitation to therapeutic strategies focused on the identification and targeting of a single mechanism. Potentially, this limitation can be bypassed by targeting collateral sensitivities associated with EMDR. Our previous work has revealed that evolving alectinib persisters exhibit a strong collateral cell-intrinsic sensitivity to a dual EGFR/HER2 inhibitor lapatinib^12^. Given the functional equivalence of EMDR and cell-intrinsic persistence and that, similar to alectinib-adapting cells, proximity to stroma is associated with partial EMT^32^, which has been linked with “stemness” and enhanced phenotypic plasticity^33–35^, we reasoned that stroma proximity might induce a similar enhancement of EGFR/HER2 dependence. Indeed, even in the absence of alectinib treatment, CAF1 CM exposure dramatically enhanced lapatinib sensitivity of H3122 and STE-1 cells in both short-term and clonogenic growth assays (**Fig. 6A, B** and **Fig. S7A, B**); the effect was observed with CM from multiple fibroblast isolates (**Fig. S7C-E**).

**Figure 6.**
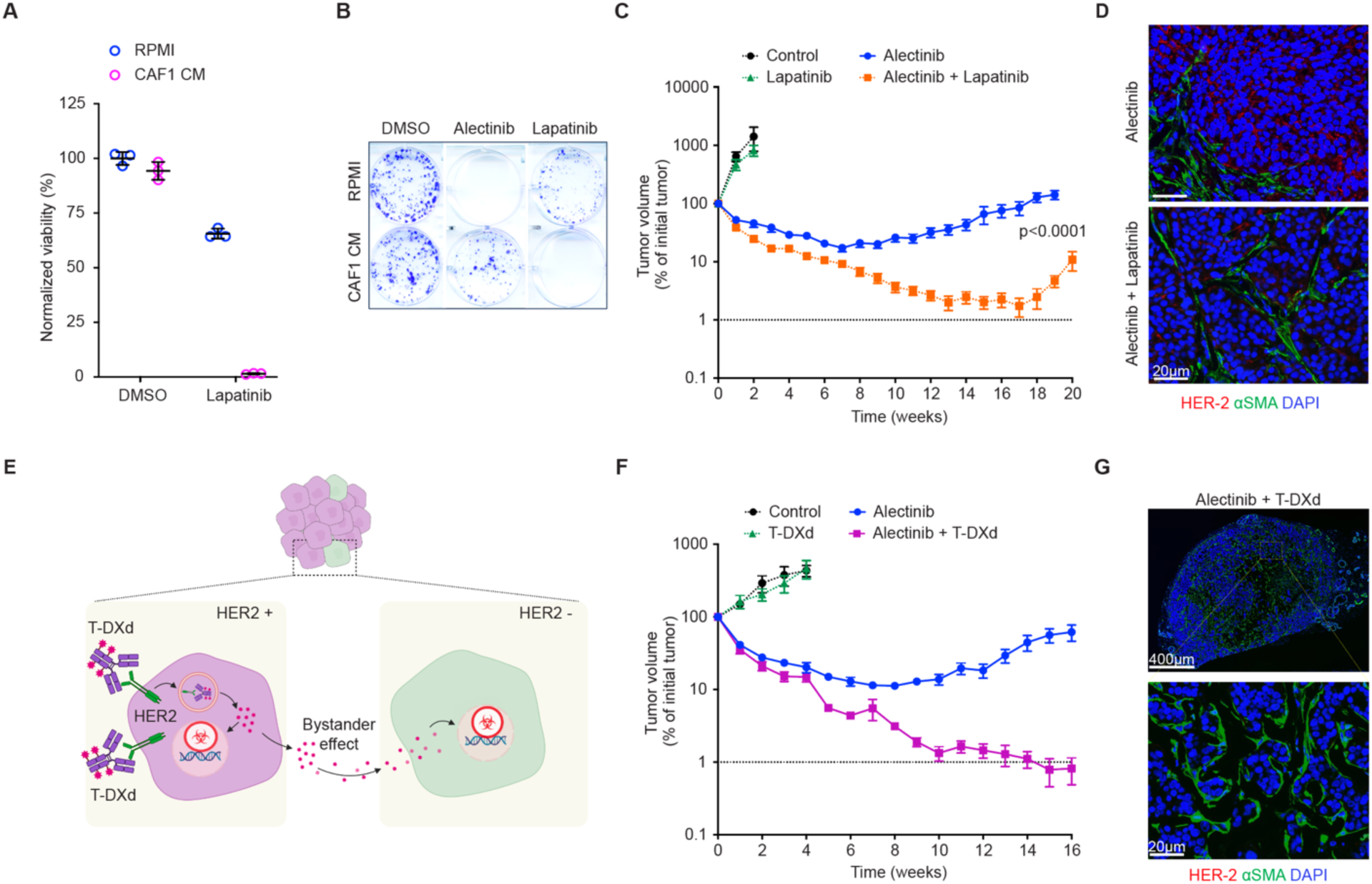
Targeting collateral sensitivity associated with stromal proximity overcomes the limitations of multifactorial resistance. **A.** Impact of CAF1 CM on sensitivity of H3122 cells to 10 µM lapatinib in 4 days CellTiter-Glo assay. **B** Impact of CAF1 CM on sensitivity of H3122 cells to 0.1 µM alectinib and 10 µM lapatinib in 10 days Crystal Violet clonogenic assay. **C**. Response dynamics of H3122 xenograft tumors treated with vehicle control (N=5), 100 mg/kg lapatinib (N=6), 25 mg/kg alectinib (N=10) and alectinib/lapatinib combination (N=10). Error bars represent SEM. **D**. Representative images of IF staining against HER-2 (red) and aSMA (green) of xenograft tumors from (**C**) at the experimental endpoint. **E**. Diagram of the experimental idea: targeting of HER2 expressing cells with the antibody-drug conjugate T-DXd should lead to elimination of the rare HER2-cells via bystander effect. **F**. Response dynamics of H3122 xenograft tumors treated with vehicle control (N=4), 10 mg/kg T-DXd (N=8), 25 mg/kg alectinib (N=8) and alectinib/T-DXd combination (N=10). **G.** Representative images of whole tumor against HER-2 (red) and aSMA (green) of xenograft tumors collected at the endpoint of the experiment shown in (**F**).

To assess the potential relevance of enhanced EGFR/HER2 dependence to H3122 cells in vivo, we examined EGFR and HER2 expression levels in the absence and presence of alectinib treatment. Whereas H3122 cells expressed detectable baseline expression of both genes, HER2 expression (**Fig. S7F**), and, to a lesser extent, EGFR expression levels (**Fig. S7G**), were further elevated after 2 weeks of alectinib treatment. Consistent with this observation and our previous report^12^, lapatinib treatment dramatically enhanced alectinib responses in H3122 xenografts, inducing stronger tumor regression compared to enhancement of individual mechanisms of EMDR (**Fig. 6C** and **Fig. S8A**). However, lapatinib co-treatment failed to overcome the ability of H3122 tumors to develop resistance and relapse. Examination of relapsed tumors indicated that most of the tumor cells have lost HER2 expression (**Fig. 6 D** and **Fig. S8B**), suggesting HER2-independent therapeutic escape.

This escape could be potentially blocked by a therapeutic modality exerting collateral damage to HER2-tumor cells. To test for this possibility, we examined the ability of HER2-targeting trastuzumab-deruxtecan (T-DXd) antibody-drug conjugate, known to have a significant bystander effect^36^, to enhance tumor sensitivity to alectinib (**Fig. 6E**). Alectinib-T-DXd combination induced a very strong regression of H3122 xenograft tumors with six out of ten tumors driven below the volumetric detectability threshold (**Fig. 6F** and **Fig. S8C**). To assess the completeness of tumor elimination, we took advantage of the mCherry fluorescent label expressed in H3122 tumor cells. Microscopic examination of the injection sites revealed a weak residual fluorescence in six out of ten sites, and subsequent histological examination validated the presence of small pockets of tumor cells. Anti-HER2 immunofluorescence staining revealed a complete loss of HER2 expression in these residual tumor cells, suggesting a stronger therapeutic bottleneck for HER2 expression then lapatinib (**Fig.S8D**).

A strong enhancement of alectinib responses was also observed in the STE-1 xenograft model. STE-1 xenograft tumors show stronger alectinib sensitivity compared to H3122, and within six weeks of treatment, multiple tumors within both alectinib (4 out of 10) and alectinib-T-DXd (8 out of 10) treated groups regressed below volumetric detectability threshold, complicating the comparison (**Fig. S8E**). To overcome this limitation, we discontinued the treatment, focusing on the relapse dynamics. Whereas discontinuation of alectinib treatment led to an immediate relapse, volumetrically detectable tumors post alectinib-T-DXd treatment continued the regression, with all of the tumors becoming undetectable, even after 11 weeks post discontinuation of treatment. Examination of the injection sites for evidence of residual disease revealed that only two out of ten sites contained small pockets of tumor cells. As with H3122 xenografts, residual STE-1 cells completely lost HER2 expression (**Fig. S8F**). Notably, the small pockets of HER2-STE-1 cells were completely encapsulated by hypertrophied fibrotic stromal tissue (**Fig. S8F**), suggesting a potential involvement of stromal reaction in suppressing tumor growth relapse. In summary, therapeutic focus on collateral sensitivities associated with persistence offers a potential solution to the dual challenge of the multifactorial nature of cell-intrinsic resistance and EMDR.

## DISCUSSION

Since therapeutic sensitivity of tumor cells is shaped by both cell-intrinsic and EMDR inputs, the ability of neoplastic populations to avoid therapeutic elimination (persistence) and to resume growth (resistance) can result from both types of inputs. However, experimental studies and clinical reports tend to attribute persistence and resistance to a single mechanism per experimental model or tumor. Historically, consideration of therapy resistance has been biased towards cell-intrinsic mechanisms. Therefore, once a cell-intrinsic mechanism is identified, it is typically considered to be sufficient to explain persistence/resistance phenotype. Our study with an experimental model of ALK+ NSCLC indicates a more complex scenario. Similar to many other models of acquired therapy resistance, cell-intrinsic mechanisms are sufficient to enable H3122 cells to survive the exposure to clinically relevant ALKi and develop resistance *in vitro*. However, proliferation of H3122 cells under ALKi within experimental tumors *in vivo* is strongly biased to the peristromal niches, even at relapse. Together with our functional validation studies, restriction of tumor cell proliferation to peristromal niches indicates the effect of EMDR (**Fig. 5E**), which can be quantitatively captured using spatial proximity metrics (**Fig. 3**). Whereas we could not fully disentangle EMDR from the effect of cell-intrinsic mechanisms, EMDR dominates the ability of H3122 tumor cells to survive therapeutic elimination, while augmenting the effects of cell-intrinsic resistance in driving tumor growth relapse. Thus, our results indicate that, at the tumor level, *in vivo* persistence and resistance can reflect a net effect of a dynamic interplay of cell intrinsic and EMDR mechanisms. Moreover, both cell-intrinsic resistance and EMDR themselves emerge through a dynamic interplay of multiple molecular mechanisms.

A large and growing body of experimental studies has documented EMDR throughout a wide range of targeted therapy contexts. Whereas the protective effect of stroma has been ascribed to multiple cell types within the tumor microenvironment, CAFs appear to be the major producer of these microenvironmental factors. Additionally, co-culture with CAFs or CAF conditioned medium has been demonstrated to reduce targeted therapy sensitivity in a wide range of therapeutic contexts^5^. Mechanistically, EMDR has been attributed to multiple stroma-produced paracrine factors, including cytokines, ECM components, metabolites, and non-coding RNAs within exosomes^5, 32^. Despite the multiplicity of documented EMDR mechanisms, individual studies tend to ascribe EMDR to a single mechanism per experimental system or tumor. Guided by this assumption of EMDR reducibility to a single mechanism, we identified HGF-cMET signaling as a pathway, predominantly responsible for CAF-mediated resistance *in vitro* (**Fig. 1**). However, our *in vivo* studies revealed a much more complex scenario: EMDR reflects the combined contribution of several growth factors (HGF, FGF and EGF family ligands) and ECM components (integrins, fibronectin and hyaluronic acid). Notably, the relative contribution of different mechanisms might depend on the response phase. For example, whereas suppression of integrin signaling through a FAKi defactinib and targeting hyaluronidase through PEGPH20 immediately enhanced alectinib sensitivity (**Fig. 5B, C**), the effect of suppression of HGF-cMET and FGF-FGFR signaling could only be detected after a significant delay (**Fig. 2C, E**; **Fig. 5C**). Importantly, our study did not exhaust the assessment of all of the potential EMDR mechanisms, uncovered in previous mechanistic studies, such as metabolic crosstalk^32^ and extracellular vehicle mediated miRNA transfer^37^. Therefore, EMDR is likely to be even more mechanistically complex, *e.g*., including the contributions of additional molecular mediators.

Given the spatial limitations of EMDR action, knowledge of its molecular mechanisms is insufficient for understanding the EMDR effects. Such mechanistic understanding would require consideration of the distance of the paracrine action, stroma/tumor ratios, and spatial distribution of the protective stroma niches. While the methodology to capture these effects has yet to be fully developed, adequate quantitative inferences can be obtained through spatial analyses of histological images using and further adapting metrics from the fields of spatial statistics and spatial ecology^38–40^ (**Fig. 3**). However, histological analyses are only capturing a single temporal snapshot. Given the temporal dimension of the acquisition of therapy resistance uncovered in recent studies including our work^12, 14, 41^, an adequate understanding of the impact of EMDR calls for consideration of both spatial and also temporal dynamics. This high complexity necessitates integrating mathematical modeling tools like spatial agent-based models^42, 43^ or compartmentalized dynamical systems^42^.

The mechanistic complexity of EMDR adds to the challenges posed by the diversity of resistance mechanisms, intratumor heterogeneity^44^ and the multifactorial nature of cell-intrinsic therapy resistance identified in recent studies^12, 14, 41^. While the development of new therapies relies on reductionistic molecular oncology studies, the ability of tumor cells to resist therapeutic elimination and the acquired therapy resistance represents a complex dynamical phenomenon, integrating the impact of multiple cell-intrinsic mechanisms, tumor heterogeneity and systemic factors (such as inflammation^45^ and variability in drug concentrations due to pharmacokinetics modifiers^46^). Consequently, the current paradigm of identifying and targeting a single mechanism at a time is unlikely to bring major clinical breakthroughs. On one hand, targeting individual mechanisms mediating EMDR or cell-intrinsic resistance can enhance the effect of the primary therapeutic agent (**Fig. 5B-D**). On the other hand, co-targeting any resistance mechanism in isolation is insufficient to prevent the ability of neoplastic populations to survive and adapt. At the same time, simultaneous co-targeting of multiple resistance mechanisms is complicated by the issue of added toxicities and the moving target nature of the tumors under treatment.

Our study highlights the potential utility of focusing on collateral vulnerabilities associated with therapy persistence. At least in the H3122 and STE-1 experimental models, both adaptive phenotypic rewiring during the acquisition of resistance to ALKi exposure to CAF produced secreted factors lead to strong enhancement in sensitivity to the dual EGFR/HER2 inhibitor lapatinib^12^ (**Fig. 6B, Fig. S7A-E**). While we do not fully understand the underpinning of this shared sensitivity, it might reflect a common denominator of partial EMT, associated with enhanced phenotypic plasticity^32, 33, 35^. Targeting this collateral sensitivity produces superior responses compared to targeting individual molecular mediators of EMDR (compare **Fig. 6C** with **Fig. 5C, D**). On the other hand, the alectinib-lapatinib combination failed to eradicate the tumors and to prevent them from adapting, likely reflecting the existence of HER2-independent therapeutic escape mechanisms. This failure prompted us to consider the use of the antibody-drug conjugate T-DXd which combines target specificity with an on-tumor bystander effect, offered by antibody-drug conjugates (**Fig. 6E**). While a subset of tumor cells still escaped eradication, the alectinib-T-DXd combination induced very strong responses, and the residual tumors were unable to re-grow after treatment discontinuation (**Fig. S8E**).

The relevance of our inferences from a relatively limited set of experimental observations that are mostly limited to a single experimental model ALK+ NCLC towards a broader range of targeted therapy contexts is yet to be investigated. However, a similarly complex interplay between cell-intrinsic resistance and EMDR has been recently reported for the development of therapy resistance in bone metastatic lesions of multiple myeloma^47^. Therefore, the attribution of therapeutic sensitivity to a combination of cell-intrinsic and EMDR inputs and the multifactorial underpinning of both types of inputs are unlikely to be limited to the H3122 model. Our results do not exclude the possibility of cases where, at any given time, therapeutic responses are dominated by either cell-intrinsic or EMDR inputs or where resistance can be attributed to a single dominant mechanism (such as gatekeeper mutations for the cell-intrinsic therapy resistance). However, the view of resistance as a dynamic spectrum of combined contribution of multiple cell intrinsic and EMDR mechanisms (with some extreme cases) offers a more generalizable and useful conceptual platform for understanding therapy resistance and developing novel therapeutic approaches. EGFR/HER2 collateral sensitivity or T-DXd-based combination therapy is unlikely to be generalizable across the broad range of ALK+ NSCLC and beyond. At the same time, identification and therapeutic targeting of shared collateral sensitivities using a therapeutic modality that addresses the issue of intratumor heterogeneity through bystander effects might offer a valuable alternative to the enumeration and targeting of individual molecular mediators of resistance.

## METHODS

### Cell culture

NSCLC cell lines H3122, STE-1, PC9, and H358 were obtained from the Lung Cancer Center of Excellence Cell Line depository at Moffitt Cancer Center. HGF, GFP and mCherry-expressing variants of H3122 cells were obtained as described ^48^. Cancer cell lines were cultured in RPMI (Gibco, ThermoFisher) supplemented with 10% fetal bovine serum (FBS), 1% penicillin/streptomycin, and 10μg/ml human recombinant insulin (Gibco) at 37°C and 5% CO_2_. All cell lines have been authenticated by short tandem repeat analysis.

Primary fibroblasts were derived from lung cancer patients’ tumors or adjacent non-cancerous lung tissue and were cultured in FB media as previously described^24^. All human tissues were collected using protocols approved by the H Lee Moffitt Cancer Center Institutional Review Board. All cancer cell lines and fibroblast isolates were routinely tested for mycoplasma contamination.

Fibroblast CM was generated by growing to ∼ 80% confluency in FB media, at which point the media was removed, the media was replaced with 10% FBS RPMI after a PBS wash. CM was collected after 48 hours, filtered (0.2 µm), aliquoted, and either used immediately or stored at −80°C. HGF concentration in the CM was determined using the human HGF ELISA Kit (Sigma-Aldrich #RAB0212) following manufacturer protocol.

For collagen coating of tissue culture plates, rat tail collagen 1 was ordered from Thermo Fischer (catalog #A1048301). Collagen 1 was further diluted to achieve 0.5 mg/ml in 0.02N glacial acetic acid, plates were coated with the standard protocol as reported by the manufacturer. Plates were then immediately used to seed cells for the experiment. Fibronectin-coated plates were ordered from Stem Cell Technologies (catalog #100-1195).

### *In vitro* viability assays

For short-term cell viability assay, 4000 cells per well were seeded in either RPMI or a 1:1 ratio of RPMI: Fibroblast CM in a 96-well plate. Treatment was initiated the next day and lasted for 3 days. Cell viability was determined using CellTiterGlo reagent (Promega) following the manufacturer’s recommended protocol. The luminescence signal was determined using a GloMax luminometer (Promega). For the analysis, the luminescence signal from the empty wells was subtracted from each reading as the background, and % cell viability was determined relative to DMSO-treated cells cultured in RPMI.

For cell viability in the co-culture setting, nuclear GFP labeled H3122 tumor cells were seeded with unlabeled fibroblasts in a 1:1 ratio along with monoculture controls, to match the total number of cells in the co-culture setting. The live cell images were acquired with the IncuCyte live cell imaging system in the green fluoresce channel (software versions 2016A/B).

For low-density long-term colony formation assays, cells were seeded in either 6-well or 24-well plates. Treatment was initiated 24 hours post seeding and lasted for 10 to 20 days. At the end point, the cells were washed with cold 1X PBS and fixed with cold methanol. Tumor cell colonies were stained with crystal violet solution (0.5% crystal violet solution in 25% methanol), followed by water wash. Plates were allowed to dry overnight and then scanned to generate the images.

### Reagents and cytokines

Alectinib, crizotinib, Lorlatinib, Ceritinib, and Cabozantinib, were purchased from Fisher Scientific. Capmatinib, Sotorasib, and ARS-1620 were purchased from SelleckChem. Stock solutions of Brigatinib, Gefitinib, and Erlotinib were received as a kind gift from Uwe Rix lab at H Lee Moffitt Cancer Center. All drugs were dissolved in DMSO, aliquoted, and stored at −20°C. For the mouse studies, pharmaceutical grade alectinib, alpelisib, pirfenidone and T-DXd were received from the Lung Cancer Center of Excellence at Moffitt Cancer Center. Infigratinib, Galunisertib and Lapatinib was purchased from AstaTech. Capmatinib and Erdafitinib were purchased from MedChemExpress. Defactinib was purchased from Selleckchem.

The human HGF Antibody was purchased from R&D systems (AB-294-NA); murine HGF recombinant protein was purchased from Sino Biological (50038-MNAH-20). EGF (GMP100-15), HB-EGF (100-47), BTC (100-50), EREG (100-04), TGF⍺ (100-16A), AREG (100-55B), BMP4 (GMP120-05ET), BMP6 (120-06), BMP7 (120-03P), TGFβ2 (100-35B), FGF1 (100-17A), FGF2 (100-18C), FGF7 (100-19), FGF10 (100-26), FGF9 (100-23), FGF18 (100-28), IL6 (200-06), IL11 (200-11), Activin A (120-14P) as well as murine growth factors & cytokines - IL11 (220-11), IL6 (216-16), FGF1 (450-33A), FGF2 (450-33), FGF7 (450-60), FGF10 (450-61), FGF9 (450-30), BTC (315-21), AREG (315-36), EGF (315-09) were purchased from PeproTech. Human recombinant NRG1 was purchased from BioLegend. We reconstituted these growth factors & cytokines in sterile 0.1% bovine serum albumin/ phosphate-buffered saline (FGF1-5mM sodium phosphate and FGF2 - 5mM sodium chloride) and were stored at −20°C.

### Animal studies

Xenograft studies were performed by subcutaneous bilateral implantation of 5E6 tumor cells/injection suspended 100 µl of 1: 1 RPMI / BME type 3 (R&D Systems #36-320-1002P). 4 to 6-week-old NOD-scid IL2Rgnull (NSG) or the NSG derivative NSG^hHGF^ (https://www.jax.org/strain/014553) recipient mice of both sexes. The animals were produced at the institute with breeders purchased from Jackson Laboratory. Treatment was initialized 3 weeks post implantation. Small molecule inhibitors were administered via daily oral gavages, 100 µl per dose. Alectinib and pirfenidone were dissolved in water. Capmatinib, alpelisib, erdafitinib, defactinib, galunisertinib and lapatinib were suspended in MCT buffer (0.25% of methyl cellulose and 0.05% of Tween 80 in water). PEGPH20 was intravenously administered at 0.1 mg/kg twice a week. T-DXd was administered via intraperitoneal injections every 2 weeks.

Tumor diameters were recorded weekly using electronic calipers; tumor volumes were calculated assuming spherical-shaped tumors. 30-45 minutes prior to euthanasia, the animals were injected IP with 10 mg/ml BrdU dissolved in 1X PBS. The weights of excised tumors were recorded, and tumors were fixed in 10% formalin solution and/or snap-frozen (in liquid nitrogen) for further analyses. All the xenograft studies were performed per the approved procedures of IACUC protocol #IS00005557 of the H. Lee Moffitt Cancer Center. Animals were maintained under AAALAC-accredited specific pathogen-free housing vivarium and veterinary supervision following standard guidelines for temperature and humidity, with a 12-hour light/12-hour dark cycle.

### Histological analyses

Formalin-fixed paraffin-embedded (FFPE) xenograft tumors were cut into 5-µm sections and mounted on glass slides. FFPE sections were deparaffinized and rehydrated, followed by heat-induced antigen retrieval in citrate buffer (pH 6.2). For immune histochemical staining, slides were incubated with 3% hydrogen peroxide in methanol followed by 10% goat serum blocking solution in 1X PBS for 10 min at room temperature. For BrdU staining, slides were incubated with primary anti-BrdU antibody (Roche,1117037600, 1:100) for 1 hour followed by the incubation with an anti-mouse biotinylated antibody (Vector Labs, BA-9200, 1:100) for 45 mins. For the detection of hyaluronic acid, slides were incubated with 4µg/ml biotinylated HABP (Sigma-Aldrich, catalog #385911) for 2.5 hours at room temperature. Detection of staining was performed with Vectastain ABC peroxidase reagent (Vector Laboratories, PK-6100) following the recommended protocol, using DAB as a colorimetric substrate and hematoxylin counterstain. Tissue collagen was detected using Masson Trichrome stain kit (Thermo Fisher, NC9752152) following manufacturer protocol. Following the staining, covering glass was mounted using permount^TM^ mounting media (Thermo Fisher, SP15-100). BrdU, HABP and Mason Trichrome stained sections were scanned with Aperio Scan Scope XT Slide Scanner (Leica).

For immunofluorescence staining, deparaffinized and rehydrated FFPE unstained tissue samples were heat incubated with antigen retrieval buffer (pH9 or pH6.2). Post one hour of incubation with blocking reagent (VectorLabs #S6000-100), tissues were incubated with primary antibodies against HER-2 (cell signaling, #4290T, 1:200) or EGFR (cell signaling, #4267, 1:50) and αSMA (Agilent, M085129, 1:100) overnight at +4C. Secondary antibody staining was performed using Alexa Fluor 488, goat anti-mouse IgG, (Invitrogen # A32723,1:1000), and Alexa Fluor 594, goat anti-rabbit IgG, (Invitrogen # A11012, 1:500) followed by nuclear staining with DAPI (Sigma-Aldrich #D9542) (1:10,000 in PBS) for 10 mins at room temperature. Prior to cover slip mounting, slides were treated with Vector Trueview autofluorescence quenching kit (VectorLabs, #SP-8400). VECTASHIELD (VectorLabs #H-1200) medium was used for slide mounting. Immunofluorescence images were acquired using PhenoCycler-Fusion system (Akoya Biosciences).

### Tissue segmentation and spatial statistics analyses

Tissue segmentation was performed using the AI-assisted digital pathology platform Aiforia version 5.2 as described in^36^. Tissues were segmented into BrdU+/- tumor cells, stroma, and necrotic tissue. The stromal areas identified by counterstaining were segmented as polygons separating cellular and acellular-ECM components. Accuracy of tissue segmentation was vetted by American Board of Pathology certified pathologist.

Spatial statistics analyses were performed using R 4.1.2. Point patterns were extracted from the Aiforia-segmented histological images as described previously^36^. Euclidian distance to the nearest stroma pixel was determined for each of BrdU+ tumor cell. Cumulative density functions of the distributions were used to perform comparison between BrdU+ and BrdU- tumor cells using the Kolgomorov-Smirnoff (KS) test. To account for the variability in tumor/stroma distribution, each sample was compared to an *in silico* random control, which retained spatial information on each of the tumor cell and stromal pixel, but BrdU+ status was randomly redistributed among tumor cells.

### RNA-seq analyses

Snap-frozen xenograft tumor tissues were immersed in liquid nitrogen and physically disrupted with a mortar pestle. Total RNA was isolated using TRIzol reagent (Invitrogen, 15596026) according to the manufacturer’s protocol. The concentration, quality, and integrity for total RNA was measured using NanoDrop and TapeStation system. cDNA library preparation and quality assessment were accomplished using Novogene’s library guidelines. NovaSeq 6000 PE150 was used as a sequencing platform. Reads were mapped to the human - GRCh38 reference genome or murine - GRCm38/mm10 reference genome. Normalized reads were obtained from DeSeq2, and principal component analysis (PCA) was performed for Hg38 and mm10, to study transcriptomic changes in human tumor cells and murine stromal component of the tumor respectively.

To compare gene expression levels across baseline, remission, and relapse, we selected a list of significant genes per sample, using the DESeq2 R package^49^. We filtered genes that have a fold change > |2| and false discovery rate (FDR) < 0.05. A ranked gene list was then created based on the genes’ log2 fold change values. Next, we generated cluster maps for Hg38 and mm10 using these DEGs. The clusters so obtained on differential expression per sample indicate genes with similar expression patterns. After clustering, we calculated the z-scores for each row (each gene) and plot these instead of the log normalized expression values; this ensured the expression patterns that we want to visualize were highlighted and not confounded by the normalized expression values. Red color indicates genes with high expression levels, and blue color indicates genes with low expression levels. The cluster maps reinforce the presence of DEGs unique to each sample, and the potential effects of treatment on the DEGs.

Gene set enrichment analysis for the mouse and human mapped genes was performed, using the KEGG7 database^50^ (www.kegg.jp) and RNAlysis software^51^ to identify the biological pathways that are significantly associated with each sample, and to assess the influence of treatment. We considered KEGG terms with p-adj < 0.05 as significant enrichment. To visualize the differential expressions of KEGG pathways across samples, we generated pathway heatmaps using the NBBt-test R package^52^.

## Statistical analyses

All statistical analyses for in vitro as well as in vivo experimental data were performed using R or GraphPad Prism software.

## Acknowledgements.

We thank Dr. Eric Lau and Karen Mann for their thoughtful comments. We thank Dr. Bradford Perez for help with acquiring primary lung tissue samples. We thank Olena Balynska for technical assistance. We thank Halozyme for generously sharing PEGPH20 reagent and the protocols. Part of the figure diagrams were created with <u>BioRender</u>.com.

---

**Supplementary Figure 1.**
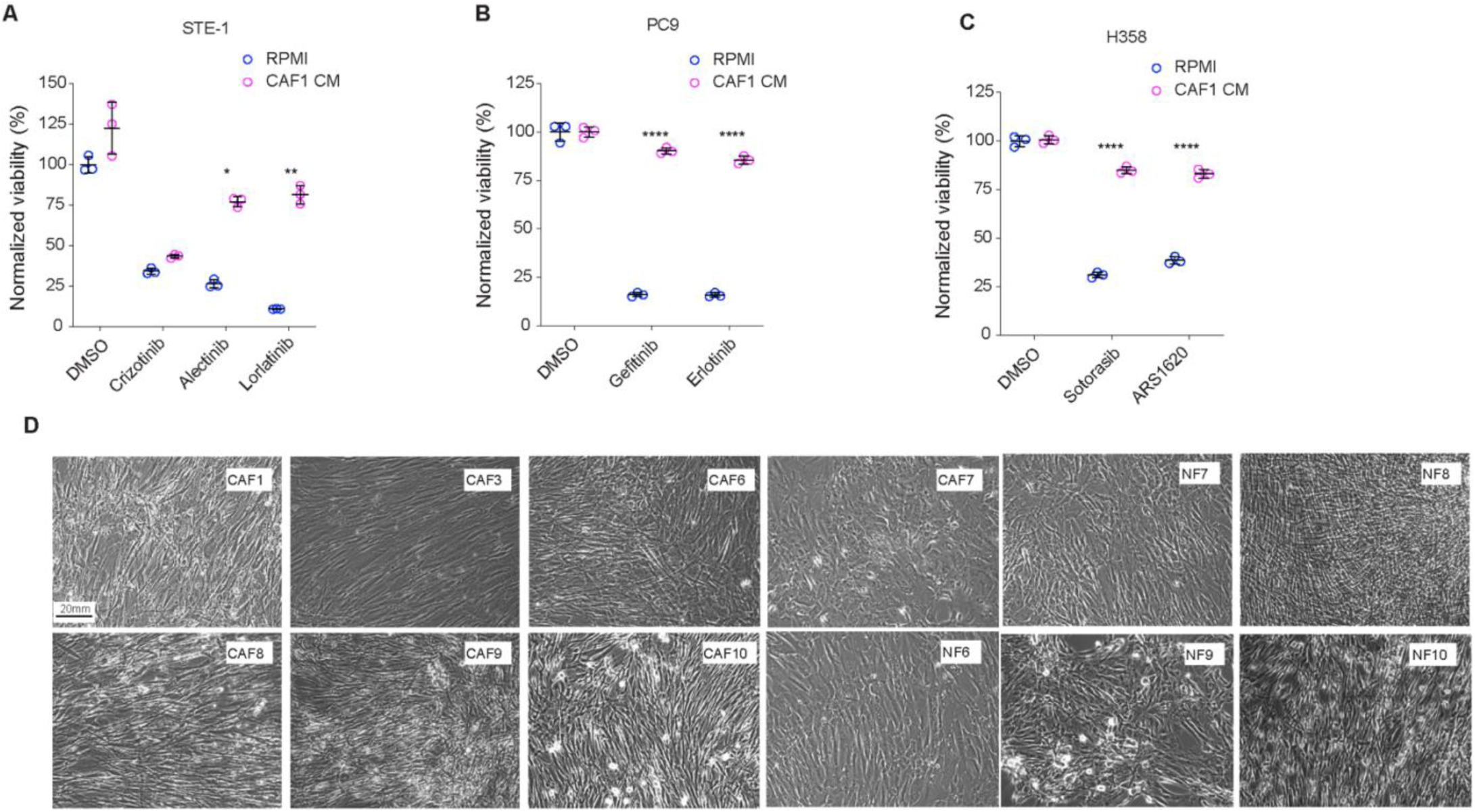
**A.** Impact of CAF CM on the viability of STE1 cells in crizotinib (0.5µM), alectinib (0.25µM), lorlatinib (0.5µM) and DMSO control. **B.** Impact of CAF CM on the viability of PC9 cells in gefitinib (0.25µM), erlotinib (0.25µM) and DMSO control **C**. Impact of CAF CM on the viability of H358 cells in sotorasib (0.05µM), ARS1620 (1µM) and DMSO control. *, **and **** refers to p<0.05, p<0.01 and p<0.0001, respectively, of the interaction term of two-way ANOVA assay, comparing the impact of CM on viability between DMSO and treatment groups. **D**. Representative microscopy images of cultures of normal and cancer-associated fibroblasts, 10x magnification.

**Supplementary Figure 2.**
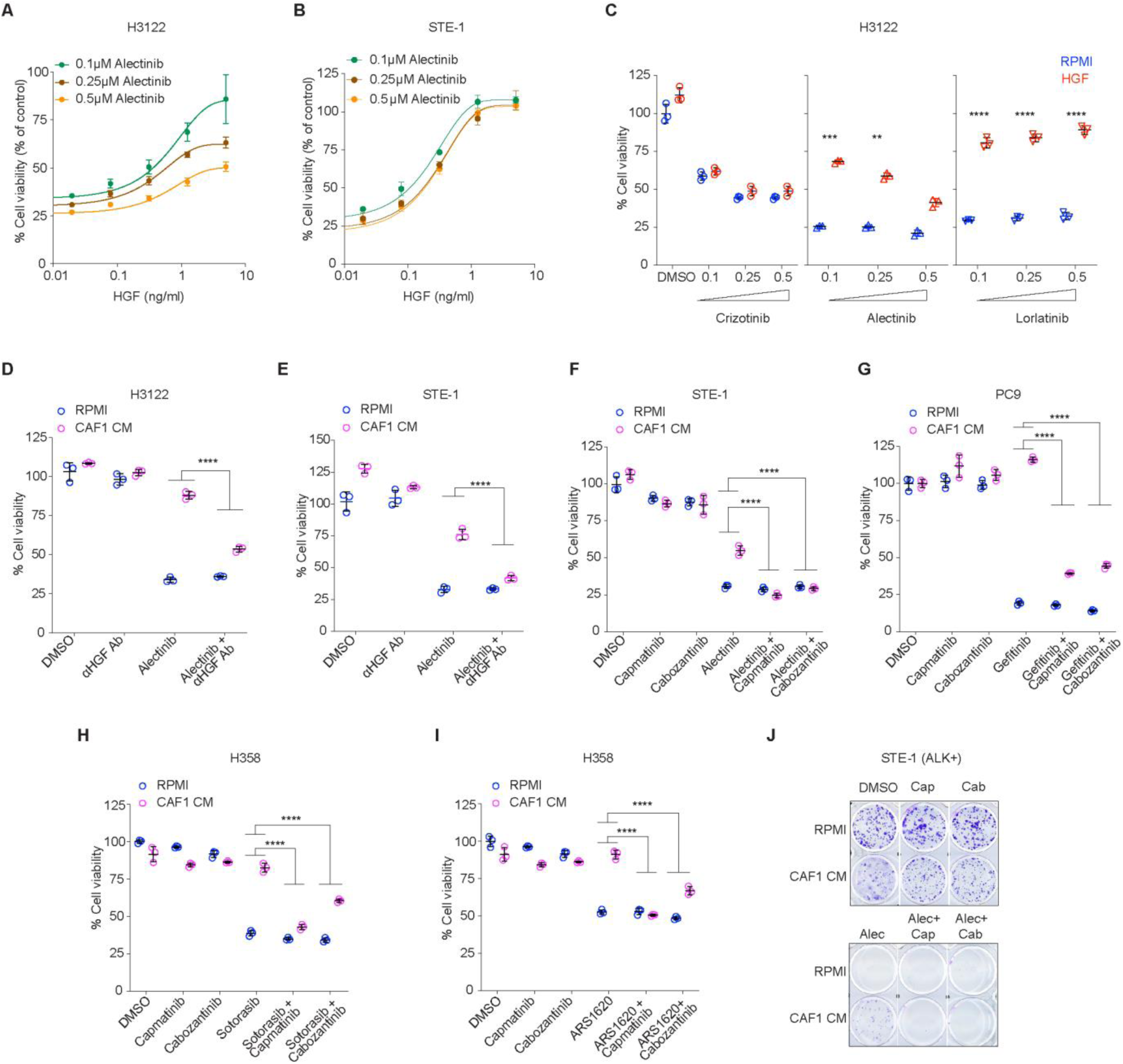
Effect of the indicated concentrations of HGF on the viability of H3122 (**A**) and STE1 (**B**) cells under the indicated alectinib concentrations. **C**. Impact of HGF (2ng/ml) on the viability of H3122 cells under the indicated concentrations of ALKi. **D**, **E**. Impact of HGF neutralizing antibody (2.5µg/ml) on the viability of H3122 (**D**) and STE1 (**E**) cells in 0.1µM of alectinib. **F-I**. Impact of cMET inhibitors cabozantinib (0.2µM) and capmatinib (0.2µM) on the sensitivity of STE-1 cells to 0.1µM alectinib (**F**), PC9 cells to 0.25µM gefitinib (**G**), H358 cells to 0.05µM sotorasib (**H**) and 1µM ARS1620 (**I**). **, ***, and **** indicate p<0.01, p<0.001, and p<0.0001 of the interaction term of 2-way ANOVA between the indicated groups. J. Microscopy images of the crystal violet stain of the 10-day culture of STE1 cells under 0.2µM capmatinib, 0.2µM cabozantinib, 0.1µM alectinib, cMETi/alectinib combinations and DMSO control.

**Supplementary Figure 3.**
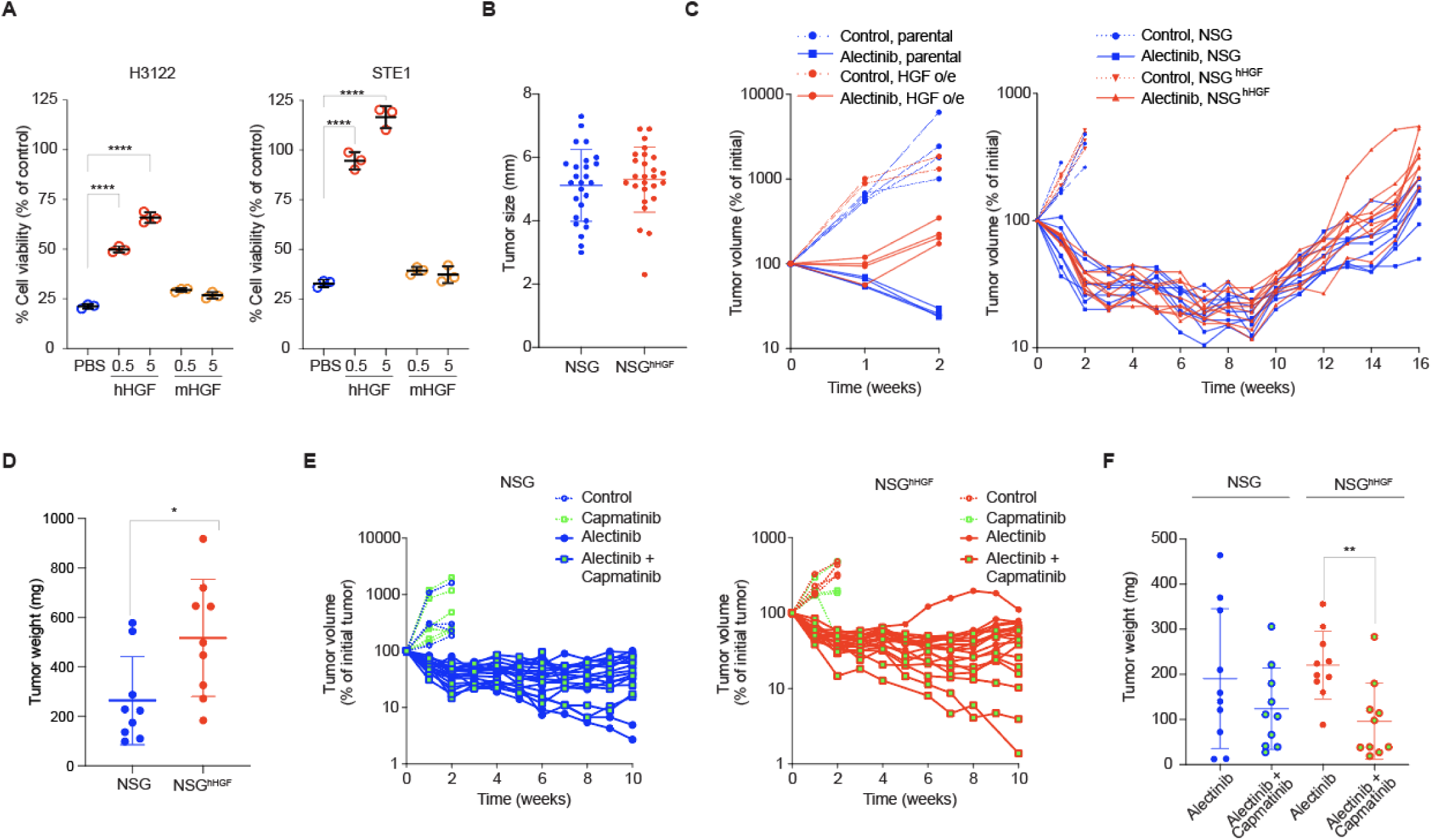
**A**. Impact of the indicated concentrations of human and murine HGF on the viability of H3122 and STE1 cells under 0.1µM alectinib. **B**. Tumor sizes (diameters) of xenograft tumors in NSG and NSG^hHGF^ hosts pre-treatment. Each dot represents an individual tumor. **C**. Volumetric traces of individual parental and HGF expressing H3122 xenograft tumors treated with 25 mg/kg alectinib or vehicle control. **C**) Volumetric traces of individual pH3122 xenograft tumors in NSG and NSG^hHGF^ hosts treated with 25 mg/kg alectinib or vehicle control. **D**. Final tumor weights from the experiment depicted in **C**. **E.** Volumetric data of individual H3122 xenograft tumors treated with 40mg/kg capmatinib, 25 mg/kg alectinib or vehicle control in NSG (left panel) or NSG^hHGF^ (right panel) hosts. **F**. Final tumor weights from the experiment depicted in **E.**

**Supplementary Figure 4.**
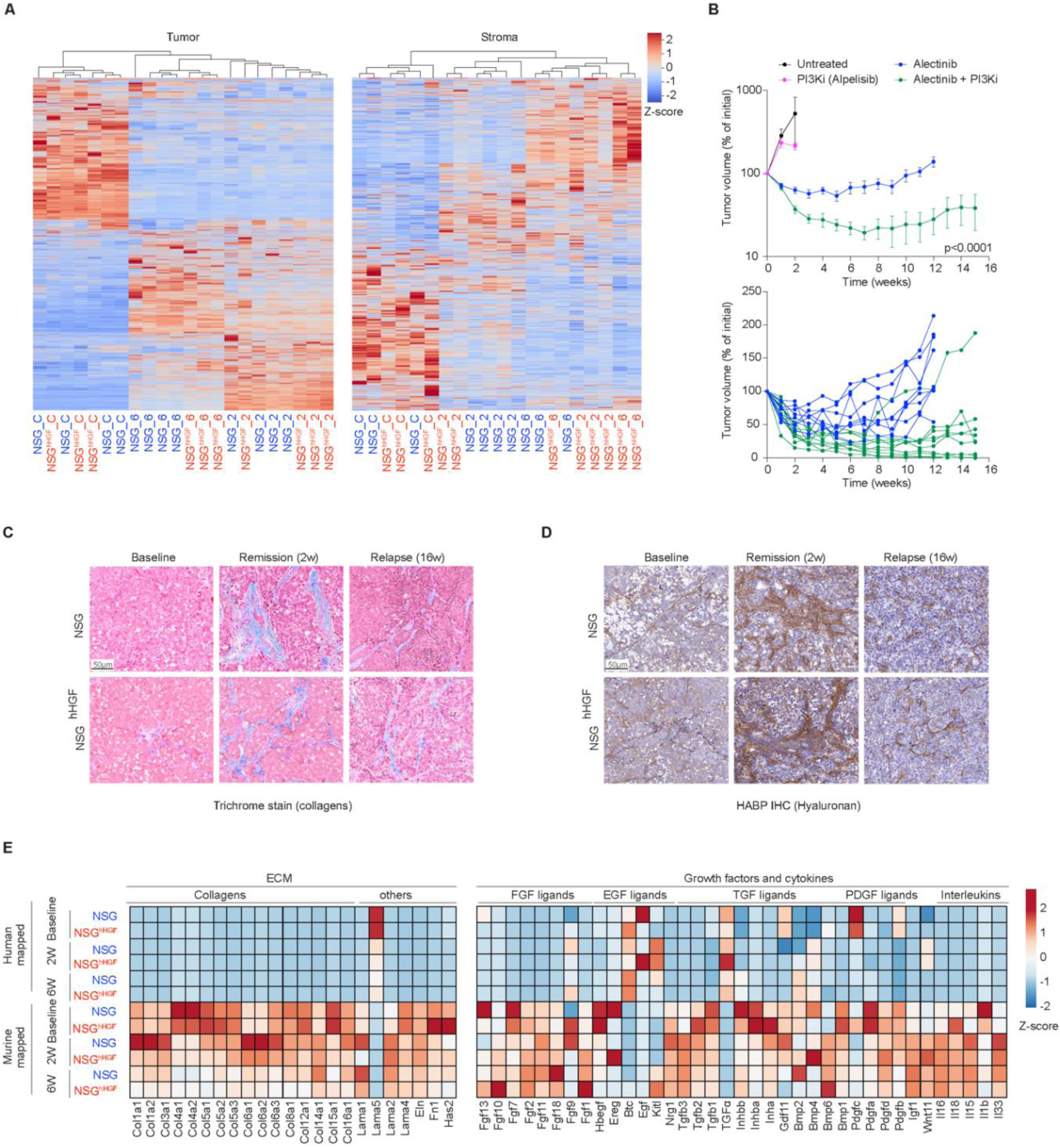
**A**. Non-supervised hierarchical clustering of the gene expression heatmaps of differentially expressed genes in the indicated samples. **B**. Volumetric responses of average (upper panel) and individual tumor traces (lower panel) of H3122 xenograft tumors in NSG hosts treated with vehicle control (N=4), 25mg/kg alpelisib (N=6), 25 mg/kg alectinib (N=8) and alectinib/alpelisib combination (N=10). Error bars represent SEM. P values represents the significance of the interaction term of 2-way ANOVA analyses between the alectinib and alectinib/alpelisib combination treated group. **C**. Representative images of Mason Trichrome staining of the indicated tumors. **D**. Representative images of the HABP IHC staining of the indicated tumors. **E**. Heatmaps of expression of the indicated genes in the indicated experimental tumors.

**Supplementary Figure 5.**
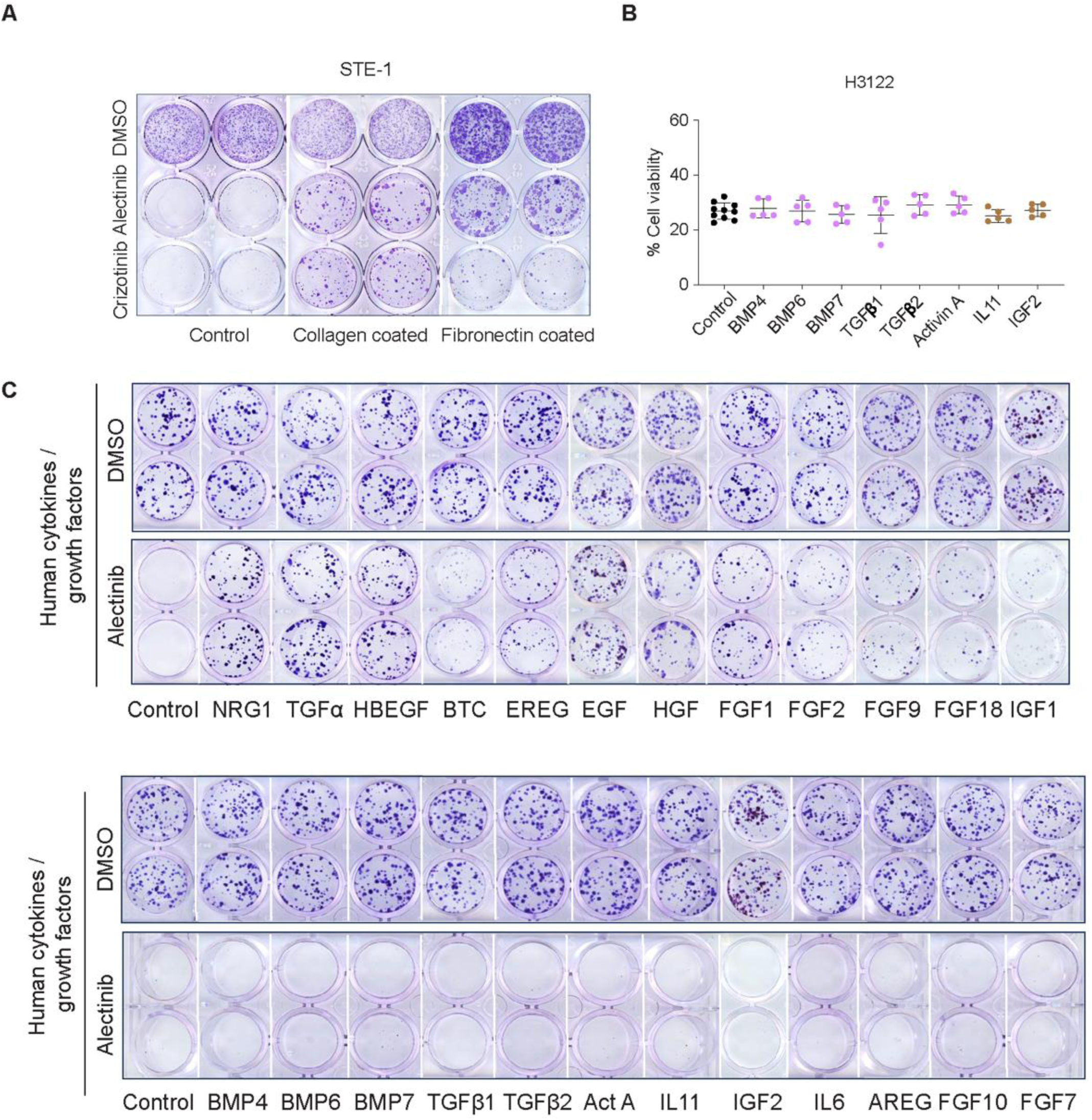
**A**. Images of crystal violet staining of the clonogenic assay of STE1 cells cultured for 20 days in regular, collagen, or fibronectin-coated plates in the presence of DMSO control, 0.1µM alectinib, and 0.5µM crizotinib. **B**. Cell viability assay for H3122 cells grown in the presence of the indicated factors (50 ng/ml). **C**. Images of crystal violet staining of the 10 days culture of H3122 cells in the presence of the indicated growth factors (50 ng/ml).

**Supplementary Figure 6.**
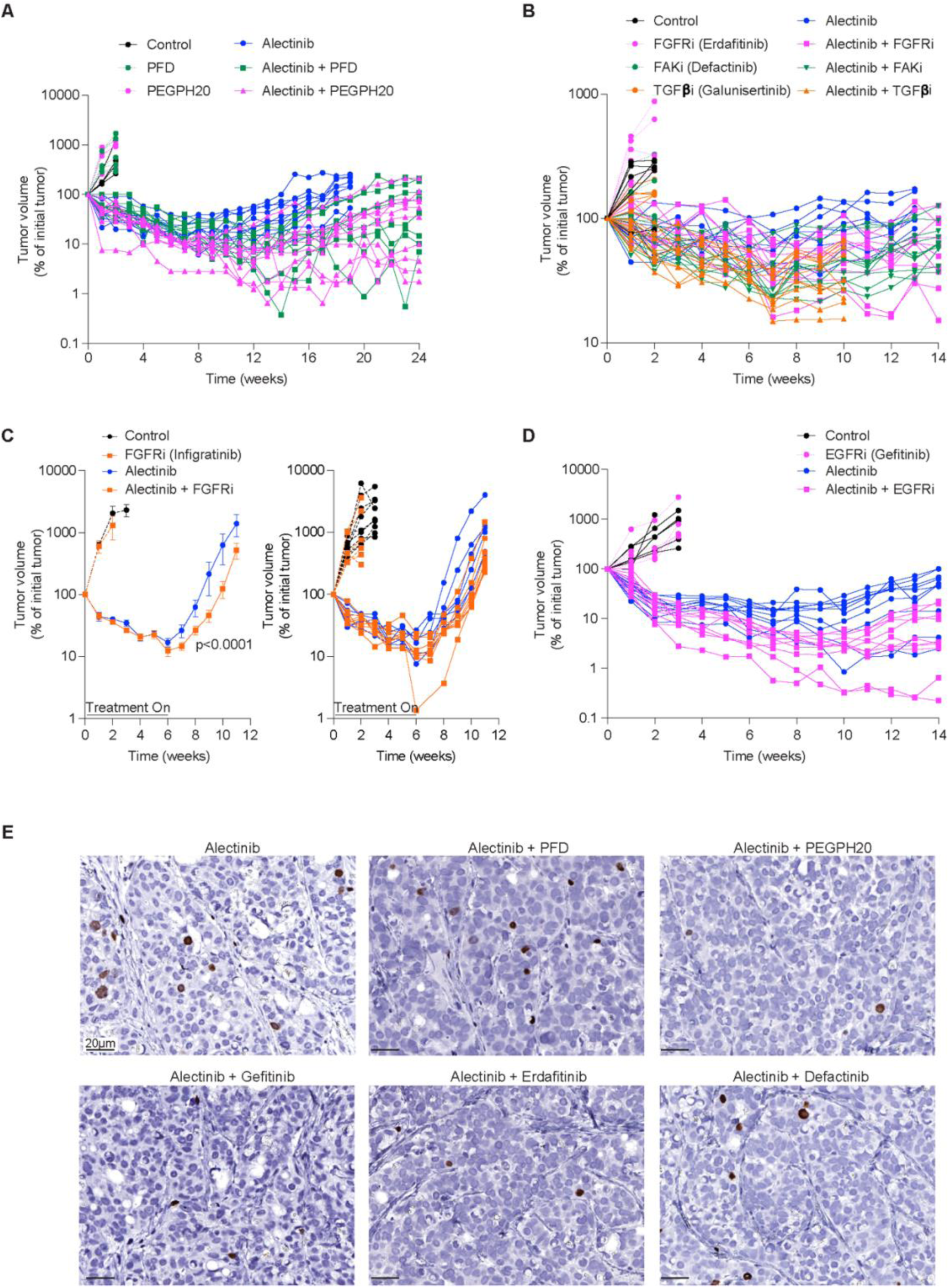
**A.** Volumetric traces of individual H3122 xenograft tumors treated with vehicle control, 900 mg/kg pirfenidone, 0.1mg/kg PEGPH, 25 mg/kg alectinib, alectinib/pirfenidone and alectinib/PEGPH20 combinations. **B**. Volumetric traces of individual H3122 xenograft tumors treated with vehicle control, 20 mg/kg erdafitinib, 50 mg/kg defactinib, 75 mg/kg galunisertib, 20 mg/kg alectinib and the indicated combinations. **C.** Volumetric traces of averages (left) and individual tumors (right) of H3122 xenograft tumors treated with vehicle control (N=10), 30 mg/kg infigratinib (N=6), 20 mg/kg alectinib (N=6), and alectinib/infigratinib combination (N=8). All mice received treatment break post 6 weeks. Error bars represent SEM. **D.** Volumetric traces of individual tumors of H3122 xenograft tumors treated with vehicle control, 40 mg/kg gefitinib, 20 mg/kg alectinib and alectinib/gefitinib combination. **E**. Representative images of anti-BrdU IHC staining of tumors tissues from H3122 xenograft mice treated with the indicated therapies for 7 days.

**Supplementary Figure 7.**
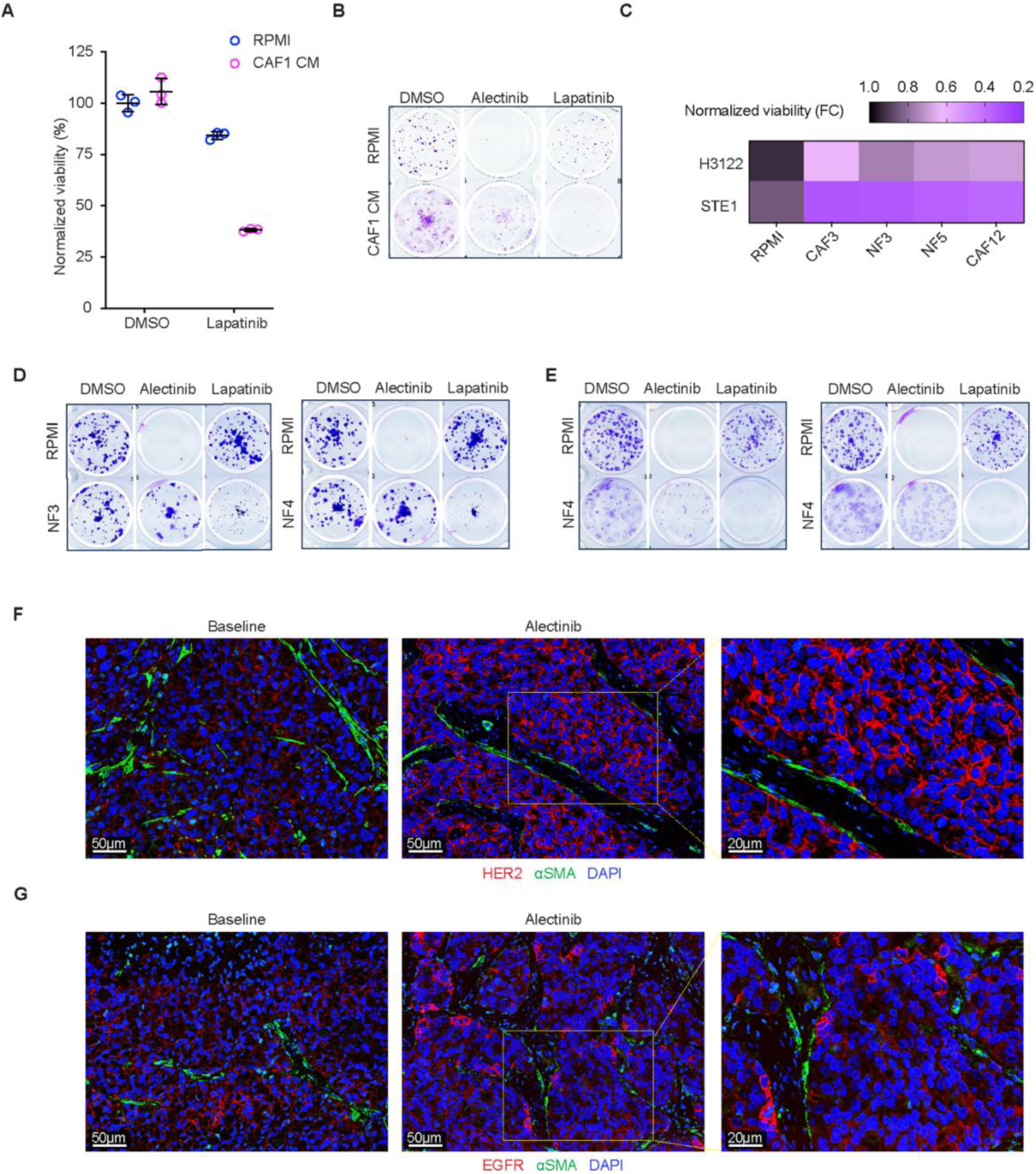
**A**. Impact of CAF1 CM on sensitivity of STE-1 cells to 10µM lapatinib in 4 days CellTiter-Glo assay. **B.** Impact of CAF1 CM on sensitivity of STE-1cells to 0.1µM alectinib and 10µM lapatinib in 10 days Crystal Violet clonogenic assay. **C.** Heatmap summary of the fibroblast CM induced lapatinib (10µM) sensitization in H3122 and STE-1 cells. The sensitivity is presented as the fold change in cell viability as compared to the RMPI media control. **D, E.** Crystal violet stain of the 10-day culture of H3122 (**D**) and STE-1 (**E**) cells in RPMI, or fibroblast CM in 0.1µM alectinib, 10µM lapatinib and DMSO control. **F, G.** Representative images of IF staining of the indicated H3122 xenograft tumors with HER-2 (red) and αSMA (green) (**F**), or EGFR (red), and αSMA (green) (**G**).

**Supplementary Figure 8.**
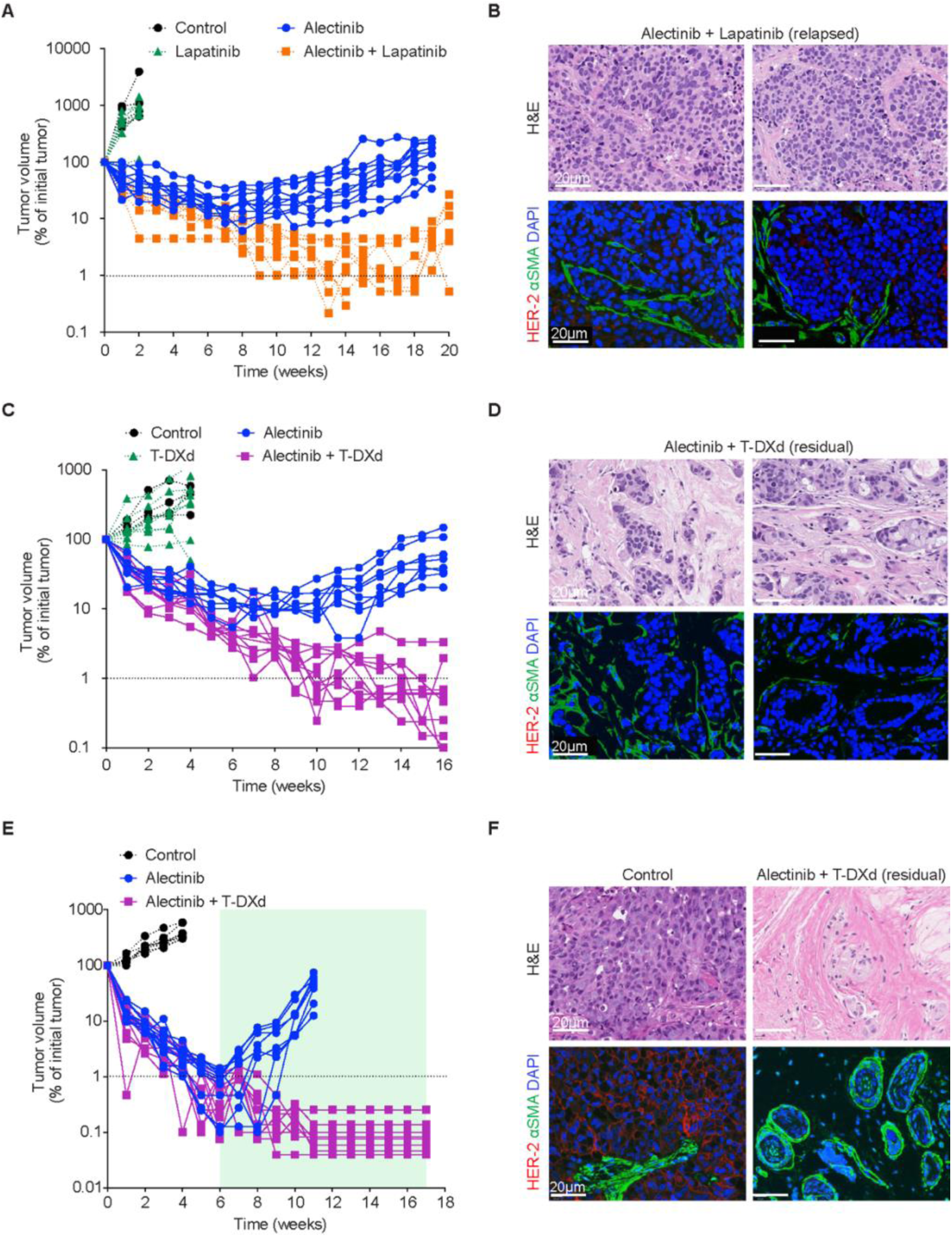
**A.** Volumetric traces of individual H3122 xenograft tumors treated with vehicle control, 100 mg/kg lapatinib, 25 mg/kg alectinib, and alectinib/lapatinib combination. **B.** Representative images of the indicated endpoint H3122 xenograft tumors from (**A**) stained with H&E (upper panel) and IF co-staining against HER-2 (red) and αSMA (green) (lower panel). **C.** Volumetric traces of individual H3122 xenograft tumors treated with vehicle control, 10 mg/kg T-DXd, 25 mg/kg alectinib, or alectinib/T-DXd combination. **_D.** Representative images of the indicated endpoint H3122 xenograft tumors stained with H&E (upper panel) and IF co-staining against HER-2 (red) and αSMA (green), DAPI (blue) (lower panel). **_E.** Volumetric traces of individual STE1 xenograft tumors treated with vehicle control, 20 mg/kg T-DXd, 12.5 mg/kg alectinib, and alectinib/T-DXd combination.**F.** Representative images of of the indicated endpoint STE1 xenograft tumors stained with H&E (upper panel) and IF and IF co-staining against HER-2 (red) and αSMA (green) (lower panel). Dotted lines in A, C and E indicate detectability threshold for volumetric measurements.

